# Sensitive multimodal profiling of native DNA by transposase-mediated single-molecule sequencing

**DOI:** 10.1101/2022.08.07.502893

**Authors:** Arjun S Nanda, Ke Wu, Sivakanthan Kasinathan, Megan S Ostrowski, Andrew S Clugston, Ansuman T Satpathy, E Alejandro Sweet-Cordero, Hani Goodarzi, Vijay Ramani

**Author notes:** These authors contributed equally to this work (Arjun S Nanda, Ke Wu, Sivakanthan Kasinathan).

## Abstract

We present SMRT-Tag: a multiplexable, PCR-free approach for constructing low-input, single-molecule Pacific Biosciences (PacBio) sequencing libraries through Tn5 transposition. As proof-of-concept, we apply SMRT-Tag to resolve human genetic and epigenetic variation in gold-standard human reference samples. SMRT-Tag requires 1-5% as much input material as existing protocols (15,000 – 50,000 human cell equivalents) and enables highly-sensitive and simultaneous detection of single nucleotide variants, small insertions / deletions, and CpG methylation comparable to the current state-of-the-art. We further combine SMRT-Tag with *in situ* adenine methyltransferase footprinting of nuclei (SAMOSA-Tag) to facilitate joint analysis of nucleosome repeat length, CTCF occupancy, and CpG methylation on individual chromatin fibers in osteosarcoma cells. SMRT-Tag promises to enable basic and clinical research by offering scalable, sensitive, and multimodal single-molecule genomic and epigenomic analyses in rare cell populations.

## MAIN

Third-generation, single-molecule long-read sequencing (SMS) technologies deliver highly accurate genomic and epigenomic readouts of kilobase (kb)- to megabase-length nucleic acid templates^1^. SMS has facilitated the characterization of previously intractable structural variants and repetitive regions^2, 3^, assembly of a gapless human genome, and high-resolution functional genomic profiling of both DNA^4–8^ and RNA^9, 10^. However, the basic and clinical applications of SMS have remained limited by the high input DNA quantities (typically >1 µg, equivalent to >150,000 human cells) required to create polymerase chain reaction (PCR)-free sequencing libraries.

Simultaneous transposition and fragmentation (*i.e.* ‘tagmentation’) using hyperactive Tn5 transposase loaded with sequencing adaptors poses an attractive solution to this problem^11^. Tagmentation serves as the basis for a variety of genomic protocols, including low-input epigenomic profiling^12–14^, cellularly-resolved monoplex^15^ and multiplex^16–18^ sequencing, highly accurate duplex sequencing^19^, and *in situ* sequencing^20^. We sought to leverage Tn5 for Pacific Biosciences (PacBio) SMS^21^ by developing <u>s</u>ingle-<u>m</u>olecule <u>r</u>eal time sequencing by tagmentation (SMRT-Tag)—an amplification-free, low-input library preparation method for simultaneously profiling the genome and epigenome on PacBio sequencers.

## RESULTS

### Multiplexed and tunable Tn5-mediated construction of PacBio libraries

SMRT-Tag (workflow shown in **Figure 1A**) relies on high molecular weight genomic DNA (gDNA) tagmentation by a triple-mutant Tn5 enzyme (hereafter referred to as Tn5), which allows concentration-dependent control of fragment size^22^. We loaded Tn5 with custom oligonucleotides composed of the hairpin PacBio adaptor and mosaic end sequences necessary for transposome assembly, and assessed the tunability of gDNA tagmentation at varying transposome concentrations and temperatures by gel electrophoresis (**Figure 1B**). This confirmed that hairpin-loaded Tn5 can effectively and tunably tagment DNA, with low temperature and low transposome concentrations favoring generation of fragments >1 kb in length.

**Figure 1:**
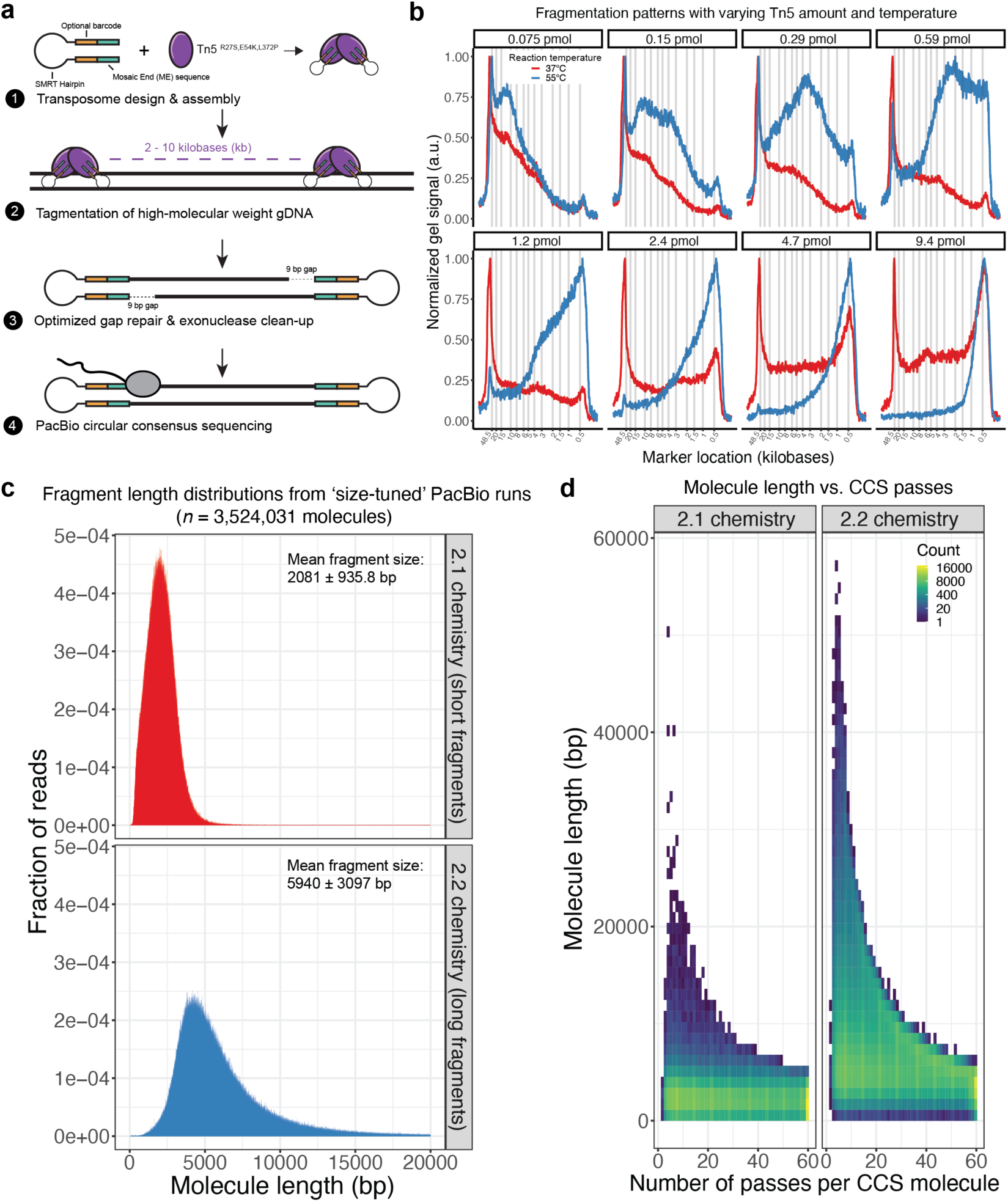
SMRT-Tag enables tunable, low-input single-molecule real time sequencing on the PacBio sequencing platform. **a.)** A schematic overview of the SMRT-Tag approach. Hairpin adaptor-loaded triple mutant Tn5 transposase is loaded and used to fragment DNA into 2 – 10 kilobase (kb) fragments. After removing Tn5 transposase, an optimized gap repair is used to fill the resulting 9 bp gaps on either side of the molecule, and an exonuclease treatment is used to purify repaired covalently closed templates, which are sequenced on the PacBio Sequel II instrument. **b.)** Hairpin-loaded transposomes can tunably fragment high-molecular weight genomic DNA (gDNA) by tuning reaction temperature and concentration. We targeted conditions that would reliably generate fragments from 2 – 10 kb in size. **c.)** Circular consensus sequencing (CCS) fragment lengths for two size-selected library preps, sequenced using size-appropriate PacBio polymerases (2.1 vs. 2.1). In red, shorter libraries sequenced using the 2.1 chemistry, and in blue, a longer library sequenced using the 2.2 chemistry. X- axis is capped at 20 kb, though 2.2 libraries exhibit a long-tailed distribution that extends past 20 kb. **d.)** Heatmap representation of molecule CCS length as a function of number of passes per CCS molecule, with log-scaled counts.

We then exhaustively-tested 62 repair conditions (**Supplementary Table 1**) to close the 9 base-pair (bp) gap created by tagmentation^23^ and allow productive PacBio sequencing. On the bases of percentage yield of DNA following exonuclease clean-up (**Supplementary Figure 1**) and fragment length estimated by analytical gel electrophoresis after tagmentation, repair, and exonuclease clean-up (**Supplementary Figure 2**), we found the highest reproducibility using two enzyme combinations: Phusion polymerase and Taq DNA ligase (‘Phusion/Taq’) and T4 DNA polymerase and Ampligase (‘T4/Ampligase’) (**Supplementary Figure 3A**). These combinations were both sensitive down to 50 ng of input gDNA, typically producing >20% total DNA yield (**Supplementary Figure 3B**). We opted to use Phusion/Taq, which worked more robustly on high quality commercial DNA samples (**Supplementary Figure 3C**), for gap repair in subsequent experiments.

To evaluate the sequencing efficiency of SMRT-Tag libraries, we tagmented 80 ng of reference-grade HG002 gDNA (equivalent to ∼15,000 cells) in a single reaction, fractionated the resulting library into two length classes using magnetic beads, and performed sequencing using PacBio’s proprietary 2.1 and 2.2 polymerases optimized for short and long templates, respectively. We generated 3,524,301 molecules over both runs (14.3 Gb total). Fragment length distributions were concordant with size-selection and polymerase choice (**Figure 1C**), with shorter mean circular consensus sequence (CCS) lengths observed with 2.1 compared to 2.2 (2081 ± 935.8 bp vs 5,940 ± 3,097 bp, mean ± standard deviation [s.d.]). Visualizing molecule length as a function of the number of individual sequencing passes per molecule (**Figure 1D**) demonstrated compatibility of these libraries with PacBio high-fidelity (‘HiFi’) sequencing, which generally requires at least 5 CCS passes per molecule to achieve 99.99% (Q20) base accuracy. We conclude that SMRT-Tag generates highly tunable libraries from low amounts of input material.

We next sought to establish multiplexability of SMRT-Tag reactions. For all sequencing reactions in this study, we used one of eight individually loaded Tn5 transposomes, each harboring a unique 8 nucleotide (nt) barcode.

To evaluate barcode demultiplexing performance, we carried out a genotype-mixing experiment using high-molecular weight gDNA isolated from previously-genotyped HG002, HG003, or HG004 human samples (**Supplementary Figure 4A**). We tagmented samples individually, pooled resulting tagged products, carried out gap-repair, and then sequenced libraries to low-depth (HG002: 0.75X; HG003: 1.39X; HG004: 1.30X). Demultiplexing using PacBio’s *lima* software demonstrated strong barcode concordance, with the vast majority of sequenced molecules harboring *lima* quality scores > 90 (mean ± s.d.: 96.76 ± 9.82; **Supplementary Figure 4B**). This suggests that transposomes neither tag previously-transposed templates following pooling, nor is there ‘barcode-hopping’ during gap repair. Finally, we assessed sample genotype mixing, with the expectation that HG003 and HG004 should cleanly separate on private genotypes, while HG002 (progeny of HG003 / HG004) should represent a mixture of both genotypes (**Supplementary Figure 4C**). As expected, while HG002 shared a moderate level (32-33%) of genotype information with HG003 and HG004, the parental samples had minimal overlap of private SNVs (0.60% HG003 vs. HG004; 0.67% HG004 vs. HG003). Taken together, these results establish the ability to parallelize and demultiplex SMRT-Tag reactions.

### SMRT-Tag accurately ascertains genomic and epigenomic variation

PacBio is thought to provide the highest genotyping accuracy across existing sequencing platforms^24^. To assess variant calling performance, we generated SMRT-Tag data from HG002 and benchmarked these data against gold-standard Genome In A Bottle (GIAB) variant callsets^25^. We sequenced HG002 to a median coverage of 11.2X across mappable genomic regions (34.24 Gb generated over 6 Sequel II flow cells) and called single nucleotide (SNVs) and insertion/deletion (indel) variants using DeepVariant^26^. Comparing SNV and indel calls from SMRT-Tag against coverage-matched data from GIAB, we observed quantitatively similar recall (0.970 vs. for 0.970 for SNVs and 0.911 vs. 0.907 for indels), precision (0.995 vs. 0.995 for SNVs and 0.955 vs. 0.949 for indels), F1 score (0.983 vs. 0.982 for SNVs and 0.932 vs. 0.928 for indels), and AUC (0.969 vs. 0.968 for SNVs and 0.902 vs. 0.897 for indels; **Figure 2A,B**). Importantly, SMRT-Tag also performed similarly in challenging genomic regions (*e.g.* segmental duplications, tandem repeats, homopolymers, and the MHC locus; **Supplementary Figure 5A**), with SMRT-Tag slightly outperforming coverage-matched GIAB in select cases, likely reflecting improvements in sequencing chemistry (F1 scores: 0.977 vs. 0.967 for SNVs and 0.912 vs. 0.905 for indels across all challenging regions). Furthermore, similarities between SMRT-Tag and GIAB variant-calling performance did not vary with respect to coverage (**Supplementary Figure 5B**), though we note that SMRT-Tag was not as effective in detecting some structural variants (data not shown), owing to shorter average fragment size in SMRT-Tag libraries compared to GIAB PacBio data.

**Figure 2:**
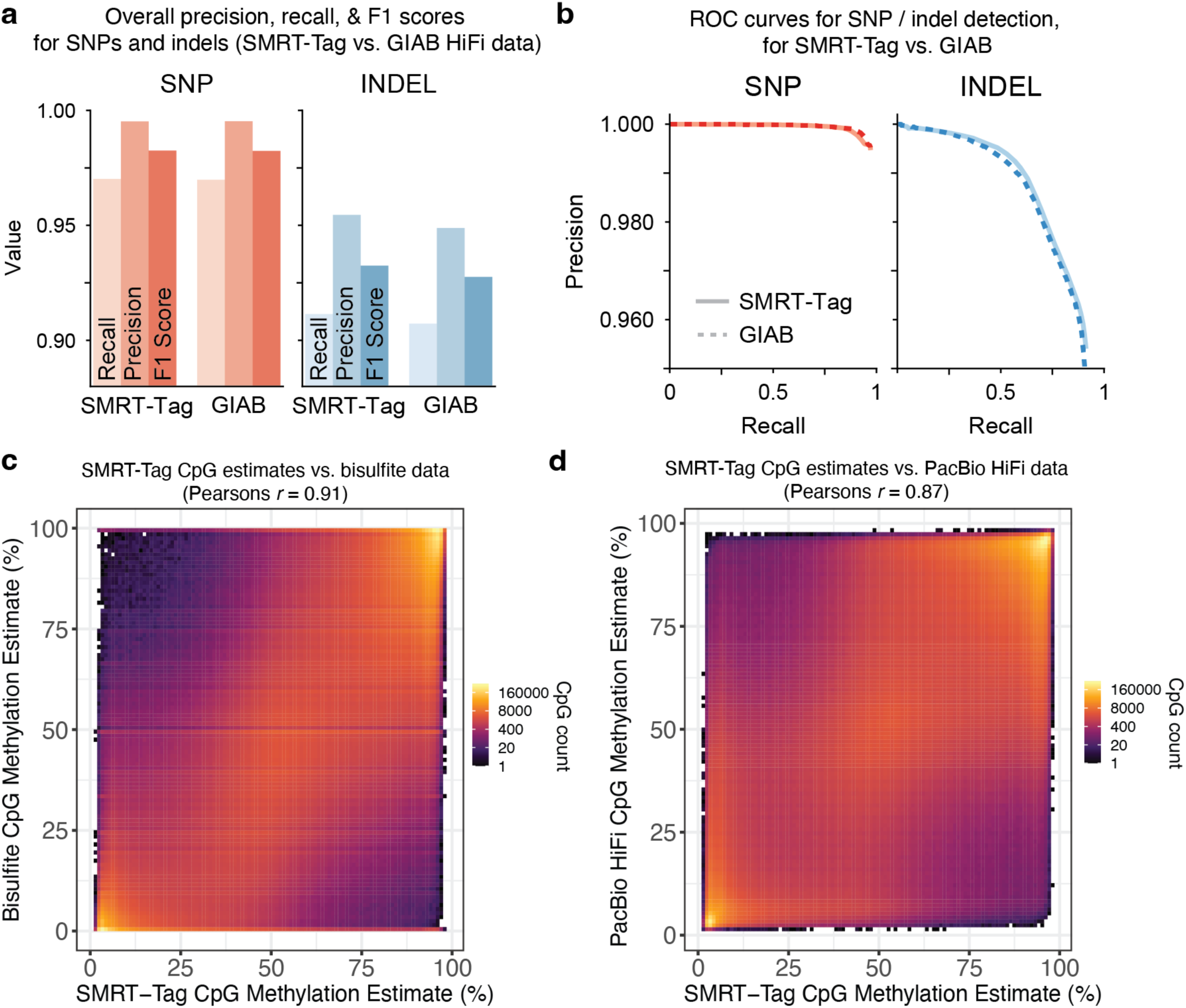
Benchmarking the genotyping and epigenotyping accuracy of SMRT-Tag reactions. **a.)** Overall precision, recall and F1 scores for DeepVariant SNP and small insertion / deletion (indel) calls in 10X SMRT- Tag HG002 data, compared against coverage-matched data from Genome In A Bottle (GIAB). **b.)** ROC curves for SNP (red) and indel (blue) detection for SMRT-Tag (solid) vs. GIAB (dashed) datasets. **c.)** Genome-wide bisulfite sequencing CpG methylation estimates for HG002 plotted against *primrose* CpG estimates from SMRT- Tag data as a binned heatmap (bin size = 1%) with log-scaled counts. Pearson’s correlation across all covered CpGs (*n* = 27,313,363 CpGs) is 0.91. **d.)** Genome-wide, downsampled GIAB Hi-Fi *primrose* methylation estimates for HG002 plotted against *primrose* CpG estimates from SMRT-Tag data as a binned heatmap (bin size = 1%) with log-scaled counts. Pearson’s correlation across all covered CpGs (*n* = 27,760,538 CpGs) is 0.87.

Sequencing native DNA on third generation sequencers offers the unique opportunity for simultaneous genotyping and epigenotyping (*i.e.* calling CpG methylation)^27^. To assess whether SMRT-Tag effectively captures HG002 CpG methylation, we ran PacBio’s *primrose* software, which uses a convolutional neural network to predict CpG modification based on real-time sequencing polymerase kinetics. We then compared genome-wide methylation estimates against publicly available gold-standard bisulfite sequencing data^28^ and against GIAB PacBio data. We observed very high correlations between per-CpG methylation calls between SMRT-Tag and benchmark bisulfite-based m^5^dC estimates (Pearson’s *r* = 0.91; **Figure 2C**), and between SMRT- Tag and downsampled GIAB PacBio data (Pearson’s *r* = 0.87; **Figure 2D**). Together, these analyses demonstrate that SMRT-Tag genotyping and epigenotyping accuracy is highly concordant with standard PacBio library preparation methods and bisulfite sequencing.

### Mapping single-fiber chromatin accessibility and CpG methylation with SAMOSA-Tag

Tn5-tagmentation of intact nuclei is the basis for ATAC-seq, a popular method for quickly and reproducibly profile bulk chromatin accessibility genome-wide^13^. To adapt SMRT-Tag to analogously assay single-molecule chromatin accessibility^4^, we developed and optimized a <u>tag</u>mentation-assisted <u>s</u>ingle-molecule <u>a</u>denine <u>m</u>ethylated <u>o</u>ligonucleosome <u>s</u>equencing <u>a</u>ssay (SAMOSA-Tag; **Figure 3A**), wherein nuclei are methylated *in situ* using the EcoGII m^6^dAase, tagmented using hairpin-loaded Tn5 under conditions optimized for ATAC-seq^29^, gap-repaired following DNA purification, and then sequenced on the PacBio Sequel II. As proof-of-concept, we applied SAMOSA-Tag to 50,000 nuclei from *MYC-*amplified OS152 human osteosarcoma cells^30^, and used a convolutional neural network hidden Markov model (CNN-HMM)^31^ to call inaccessible protein-DNA interaction ‘footprints’ from m^6^dA modifications natively detected by the sequencer. In total, across eight replicates, we sequenced 3,640,652 single molecules (7.79 Gb). Consistent with transposition of chromatin in nuclei, SAMOSA- Tag CCS length distributions displayed a characteristic oligonucleosomal banding pattern at shorter lengths (**Figure 3B**). When aligned to 5’ read ends, SAMOSA-Tag molecules further displayed periodic accessibility signal, consistent with Tn5 transposition adjacent to nucleosomal barriers (**Figure 3C**). Sizes of individual footprints corresponded with expected sizes of mono-, di-, tri-, etc. nucleosomes (**Figure 3D**). Finally, single fiber accessibility patterns could be visualized in the context of the genome, for instance at the amplified *MYC* locus (**Figure 3E**). Importantly, unlike ATAC-seq data, SAMOSA-Tag insertions were only mildly biased toward annotated transcription start sites (TSSs; **Supplementary Figure 6A**); insertions did, however, preferentially occur in the vicinity of predicted CCCTC-binding factor (CTCF) binding sites (**Supplementary Figure 6B**), consistent with blocking of Tn5 transposition by strong barrier elements. This slight insertion preference was also reflected in the overall fraction of insertions falling within TSSs and around CTCF binding sites (**Supplementary Figure 6C**; 1.51-fold enrichment above background for TSS; 1.58-fold enrichment above background for CBS), and was consistent with previously reported biases for Tn5-mediated shotgun Illumina sequencing^32^.

**Figure 3.**
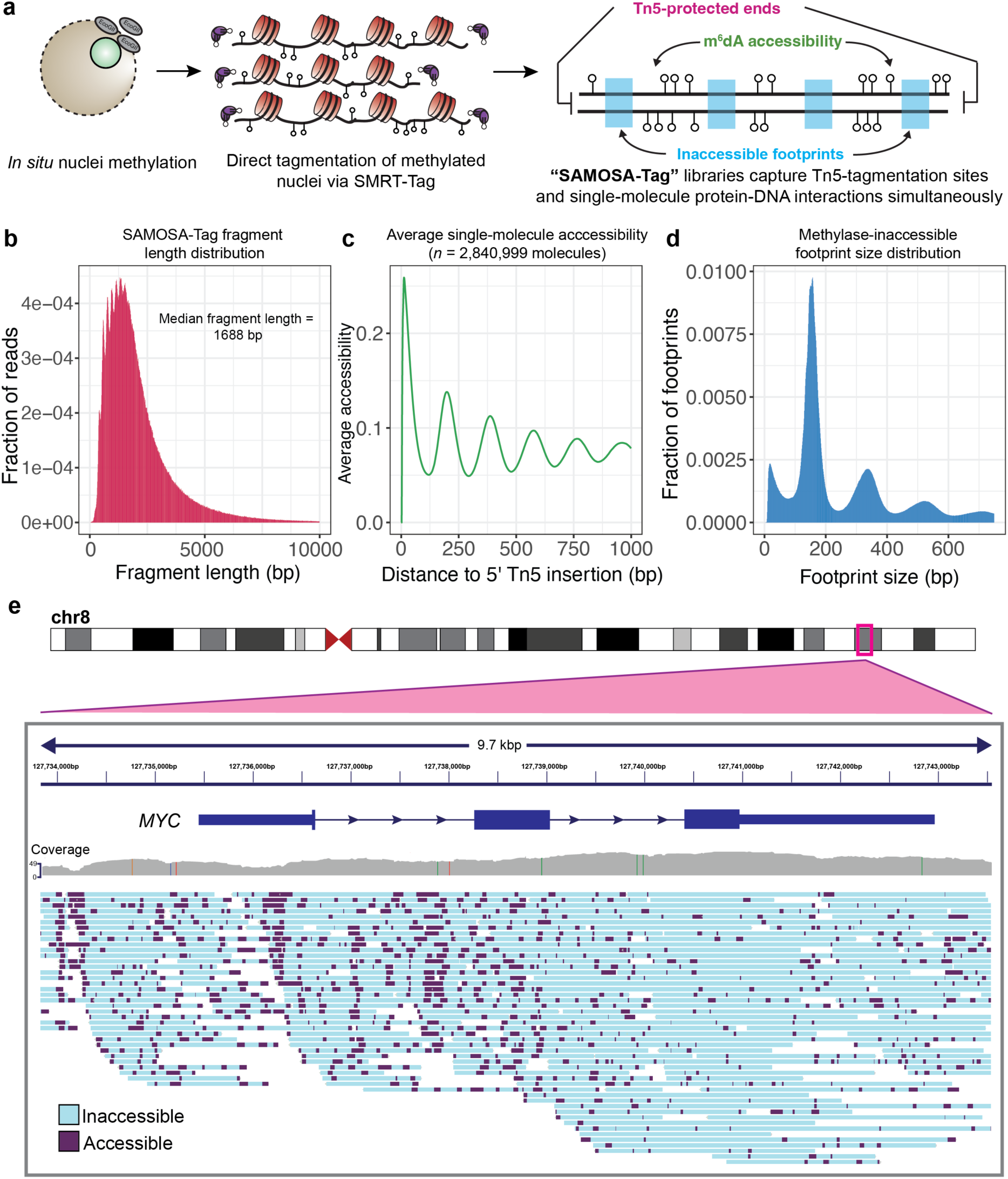
SMRT-Tag can be combined with the SAMOSA single-fiber footprinting assay to easily generate single-molecule chromatin accessibility data through direct tagmentation of adenine-methylated nuclei. **a.)** Schematic overview of the SAMOSA-Tag approach: nuclei are methylated using the nonspecific EcoGII m^6^dAase and tagmented *in situ* using SMRT-Tag. DNA is purified, gap-repaired, and sequenced on the PacBio Sequel II, resulting in molecules where ends result from Tn5 transposition, m^6^dA marks represent fiber accessibility, and computationally defined unmethylated ‘footprints’ capture protein-DNA interaction. **b.)** Fragment length distributions for SAMOSA-Tag data from the OS152 osteosarcoma cell line. **c.)** Average methylation signal from the first 1000 nt of molecules from the same dataset as **b.). d.)** Unmethylated footprint size distribution for the same dataset. **e.)** Genome browser visualization of SAMOSA-Tag data at the amplified *MYC* locus. Purple marks predicted accessible bases, while blue represents predicted inaccessible bases on individual molecules.

### Integrative measurement of CpG methylation and single-molecule chromatin accessibility

We speculated that separation of SAMOSA-Tag polymerase kinetics into separate m^6^dA and m^5^dC channels would enable simultaneous readout of DNA sequence, CpG methylation, and chromatin fiber accessibility. We first examined accessibility and CpG methylation signal surrounding predicted CTCF binding sites derived from ChIP-seq in the U2OS cell line, with the expectation that OS152 and U2OS should share a subset of binding sites. Averaged accessibility and CpG methylation signals in 750 nt windows centered at predicted CTCF motifs revealed characteristic hallmarks of CTCF binding, including positioned nucleosomes flanking the motif, decreased fiber accessibility immediately at the motif (consistent with exclusion of EcoGII by fiber-bound CTCF), and depressed CpG methylation within motifs (**Figure 4A**). To move past this fiber average, we used unbiased Leiden clustering^33^ to examine the different fiber structures that make up this pattern (example of 4 clusters shown in **Figure 4B;** cluster sizes shown in **Supplementary Figure 7**). Analysis of average CpG methylation associated with each fiber structural pattern (**Figure 4C)** revealed lowest CpG methylation in clusters displaying direct evidence of CTCF fiber binding (cluster 1; minimum unsmoothed m^5^dC/C of 0.14) and motif accessibility without bound CTCF (cluster 2), consistent with prior results^34^. Two additional analyses confirmed minimal confounding of CpG and m^6^dA methylation signals: i.) *primrose* score distributions between negative control (*i.e.* SAMOSA-Tag experiments where EcoGII methylation was omitted) and footprinted samples were concordant (**Supplementary Figure 8A**), and ii.) average CpG methylation signal surrounding predicted CTCF sites on fibers without detectable accessibility was tightly correlated with signal from fibers with observed footprints (**Supplementary Figure 8B**). The combination of *in situ* chromatin footprinting and tagmentation in SAMOSA-Tag illustrates the tractability of exploiting the inherent multimodality of SMS to jointly assay protein-DNA interactions, epigenetic modifications, and DNA sequence in a single experiment.

**Figure 4.**
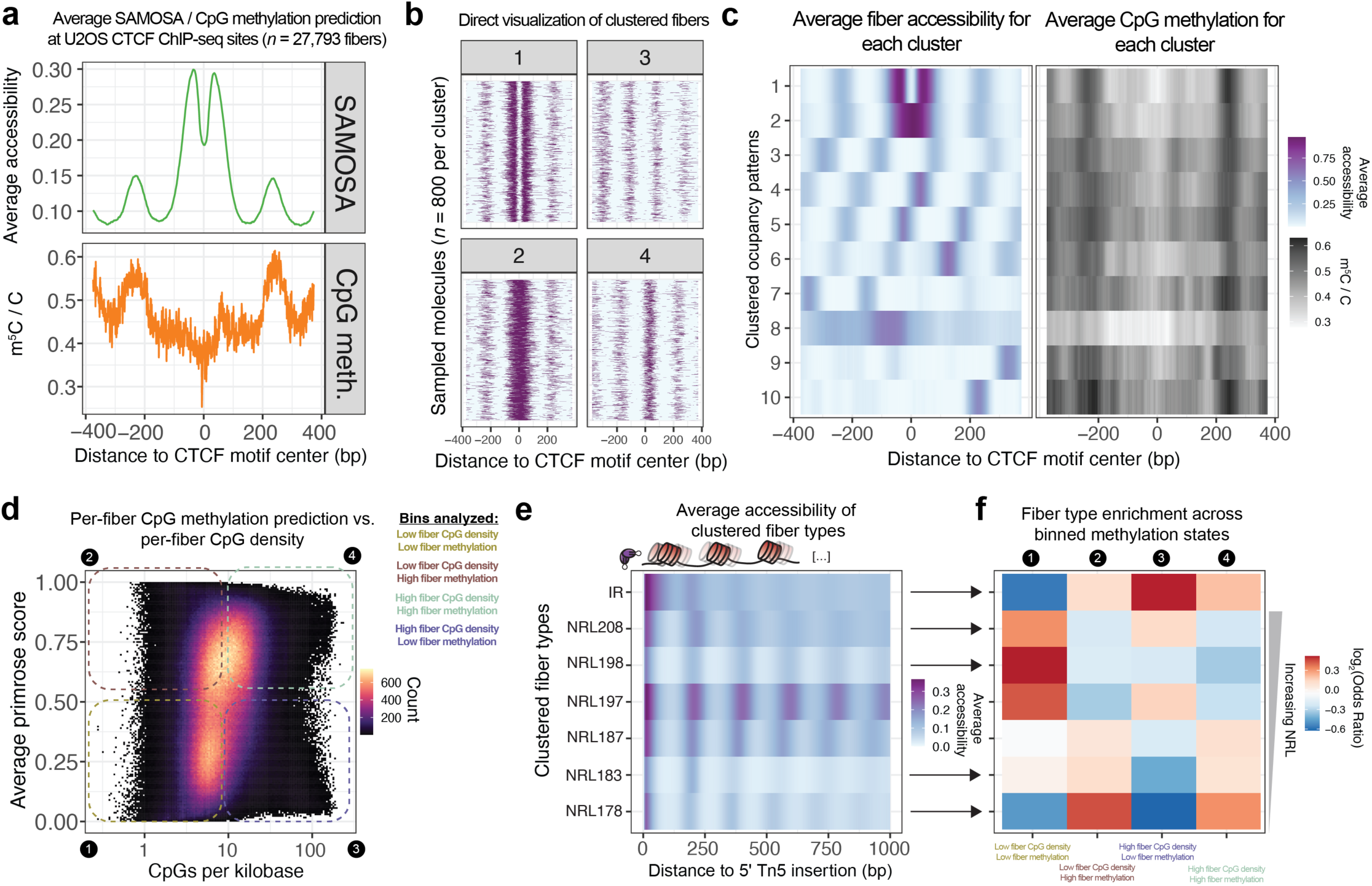
SAMOSA-Tag data can simultaneously ascertain CpG methylation state and chromatin accessibility at predicted CTCF binding sites, and can be used to study chromosome fiber structure on differentially CpG methylated fibers. **a.)** Average SAMOSA accessibility signal and CpG methylation on 27,793 footprinted fibers, centered at predicted CTCF binding sites taken from published U2OS CTCF ChIP-seq data. **b.)** Molecular visualization of individual, clustered fibers (800 molecules per cluster), reflecting different CTCF-occupied, accessible, and inaccessible fiber states, centered at predicted CTCF binding motifs. **c.)** Simultaneous visualization of average accessibility (left) and CpG methylation (right) for each of 10 clustered accessibility states surrounding CTCF motifs. Window size is 750 nt for **a.) – c.). d.)** Average *primrose* score (methylation prediction) for individual fibers as a function of number of CpG dinucleotides per kilobase on individual fibers. We binned molecules into one of four bins, depending on both CpG density and average *primrose* score. **e.)** Average accessibility of 7 different fiber types determined by performing leiden clustering on single-molecule autocorrelograms calculated from each footprinted chromatin fiber. Clusters broadly stratify the entire genome on the basis of NRL for regular fibers (ranging from 178 to 208 nt), or irregularity (cluster IR). **f.)** For the same clusters as in **e.)**, relative enrichment or depletion (calculated through Fisher’s exact test) of individual fiber types in each of the four binned states from **d.).** All tests shown here are statistically significant (*p*-values range from ∼0 to 2.41E-05).

In prior work, we demonstrated that single-fiber accessibility data could be used to cluster the genome based on nucleosome regularity and average distance between regular nucleosomes (nucleosome-repeat length, or NRL)^4, 31^. These studies relied entirely on complementary epigenomic datasets to assess how the distribution of so-called ‘fiber-types’ (*i.e.* collections of fibers with unique regularity or NRL) differed across euchromatic and heterochromatic domains. We sought to improve on these analyses by directly assessing how fiber structure varies as a function of jointly-measured single-molecule CpG content and CpG methylation. To do so, we assessed the distribution of single-molecule CpG densities, and average *primrose* methylation scores for each sequenced SAMOSA-Tag molecule in our dataset (**Figure 4D**). We then sectored these molecules into four different bins, gated on CpG density (> 10 CpG dinucleotides per kilobase), and *primrose* score (average *primrose* score > 0.5). We then (as previously^4, 31^; **Methods**) computed single-molecule autocorrelograms for each sequenced molecule at least 1 kb in length, and clustered autocorrelograms to define fiber types. Following filtering of artifactual molecules, we obtained 7 distinct clusters (**Figure 4E**; cluster sizes in **Supplementary Figure 9**), which effectively stratified the OS152 genome by NRL (clusters NRL178 – NRL208) and fiber regularity (cluster IR). Finally, using the methylation / CpG content bins, we carried out a series of enrichment tests to assess how these fibers were differentially distributed across high / low CpG content and predicted CpG methylation (**Figure 4F;** reproducibility shown in **Supplementary Figure 10**). The resulting heatmap relates domain-specific changes in fiber composition as a function of single-molecule CpG state, and we highlight two findings that suggest relevance to chromatin regulation: first, we find that high CpG content / low CpG methylation (*i.e.* likely hypomethylated CpG islands) fibers are enriched for irregular fibers (odds ratio [O.R.] for cluster IR = 1.42; *p* ∼ 0), as well as fibers with long NRLs (NRL208 O.R. = 1.09 / *p* = 4.43E-64; NRL197 O.R. = 1.11 / *p* = 1.49E-58); second, we find that high CpG content / high CpG methylation fibers (*i.e.* likely hypermethylated, CpG rich repetitive sequence) are enriched for irregular fibers (IR O.R. = 1.14 / *p* = 1.33E-139), as well as short NRL fibers (NRL172 O.R. = 1.24; *p* ∼ 0). Both results are broadly consistent with our previous *in vivo* SAMOSA observations of active promoters and heterochromatin in human K562 cells^4^ and murine embryonic stem cells (mESCs)^31^, pointing to a conserved pattern of single fiber chromosome structure within these domains. Together, these analyses demonstrate that SAMOSA-Tag can easily generate genome-wide, multi-omic single-molecule chromatin fiber accessibility data from tens of thousands of cells.

## DISCUSSION

Here, we demonstrate direct transposition for sensitively preparing multiplexed, amplification-free PacBio sequencing libraries. We apply this principle to develop two related, single-molecule native DNA sequencing approaches: i.) SMRT-Tag, which enables highly-accurate genomic variant and methyl-CpG detection comparable to gold-standard WGS and bisulfite data, and ii.) SAMOSA-Tag, which adds a third, concurrent readout of protein-DNA interactions on individual chromatin fibers using *in situ* m^6^dA footprinting. Crucially, combining tagmentation with optimized gap repair allowed the streamlined creation of PacBio libraries from 80 – 100 ng DNA (a minimum of 15,000 human cell equivalents) compared to current protocols that require > 0.5 – 5 μg DNA (a minimum of ∼200,000 human cell equivalents). We anticipate that this order-of-magnitude reduction in input requirement will remove a major obstacle to routine PacBio sequencing and empower basic and translational studies of rare cell populations.

We also address the need for functional genomic techniques capable of characterizing the breadth of nucleotide, structural, and epigenetic variation ascertained by SMS. Like ATAC-seq, SAMOSA-Tag offers a straightforward, scalable method for rapidly profiling single-fiber chromatin accessibility, CpG methylation, and genetic variation from as few as 50,000 nuclei. Our proof-of-concept raises the possibility of single-cell resolution SMS approaches for profiling native chromatin and DNA. We envision the further development of SMRT-Tag to include droplet- or combinatorial barcoding-based cellular indexing: such approaches would extend multimodal long-read analyses to the resolution of hundreds to thousands of individual cells in parallel, enabling applications ranging from somatic variant detection, to *de novo* assembly, to cell type classification.

While SMRT-Tag and SAMOSA-Tag are powerful tools, we note that both approaches have limitations. Because SMRT-Tag does not rely on PCR, optimal use of individual PacBio flow cells requires multiplexing SMRT-Tag reactions; in experiments presented here, we saturated PacBio flow cell loading by combining barcoded, repaired templates from 6 – 8 SMRT-Tag reactions. Further, owing to limited input, SMRT-Tag reactions are not compatible with pulsed field or other gel-based size-selection procedures; product size distributions in SMRT- Tag are primarily controlled by transposome concentration and bead-based cleanup protocols, and unsequenced DNA is effectively lost. This limitation is particularly important for large-scale structural variant discovery, as the abundance of long, breakpoint-spanning CCS molecules will be lower in SMRT-Tag libraries compared to gold-standard HiFi data. While we have partially addressed this by separately sequencing “both sides” of an AMPure cleanup to maximize library recovery, future work optimizing tunability of transposases will enable on- demand generation of libraries with desired fragment length distributions. Similarly, our SAMOSA-Tag protocol is limited with respect to the minimal amount of nuclei that can be processed. In experiments presented here, we were able to generate high-quality data from 50,000 footprinted nuclei; future optimizations to the SAMOSA- Tag protocol including miniaturized methylation reactions or immobilization of nuclei on activated beads^35^ could further decrease this input requirement.

More generally, SMRT-Tag and SAMOSA-Tag add to a growing series of technological innovations centered around third-generation sequencing, including Cas9-targeted sequence capture^36^, combinatorial-indexing-based plasmid reconstruction^37^, and concatenation-based isoform-resolved transcriptomics^38^. The widespread adoption of short-read genomics in basic and clinical applications was catalyzed by the development of tools that democratized sequencing. Our approaches offer similar promise for rapidly maturing third-generation sequencing technologies through scalable, sensitive, and high-fidelity telomere-to-telomere genomics and epigenomics.

## DATA AVAILABILITY

Raw sequencing data is deposited in the NCBI Sequence Read Archive under accession number PRJNA863422 and in the Gene Expression Omnibus under accession number GSE210136.

## CODE AVAILABILITY

Scripts used to perform analyses are available via GitHub at https://github.com/RamaniLab/SMRT-Tag.

## ACKNOWLEDGEMENTS

We thank members of the Ramani and Goodarzi labs, Ryan Corces (Gladstone Institutes), and R. Majzner (Stanford) for critical discussion of this work. We thank the Stanford Research Computing Center for providing computational resources that contributed to this study. This work was supported by NIH grants DP2HG012442 (V.R.), R01GM123977 (H.G.), a Pew-Stewart Scholars for Cancer Research Award (A.T.S.), R01CA243555 (E.A.S.C), and a physician-scientist development award from the Stanford Department of Pediatrics (S.K.).

## AUTHOR CONTRIBUTIONS

S.K. and V.R. conceptualized the approach. A.S.N., K.W., S.K., V.R., and H.G. designed the study. A.S.N. and K.W. performed the experiments. M.S.O. assisted with preparation of sequencing libraries. A.S.C. and E.A.S.C. assisted with the osteosarcoma experiments. V.R., S.K., A.S.N., and K.W. analyzed data with support from A.T.S. V.R., S.K., K.W., and A.S.N. wrote the manuscript with input from all authors. V.R., H.G., and S.K. supervised the work.

## METHODS

### Cell lines and cell culture

OS152 cells were obtained from Alejandro Sweet-Cordero Laboratory at UCSF and cultured in standard 1x DMEM supplemented with 10% Fetal Bovine Serum and 1% 100x Penicillin-Streptomycin-Glutamine.

### Assembly of SMRT-Tag transposome complexes

#### Annealing SMRT-Tag Adaptors

HPLC-purified unique SMRT-Tag adaptors were purchased from IDT (Coralville, IA) and normalized to 100µM in RNAse-free water. Adaptors were subsequently diluted to 20µM in 1x Annealing Buffer (10mM Tris-HCl pH 7.5 and 100mM NaCl), annealed via thermocycler (95°C 5 min, RT 30 mins, 4°C hold), and rapidly cooled to −20°C for long-term storage.

#### Loading Tn5 transposases with SMRT-Tag adaptors

Purified H-Tn5^R27S,E54K,L372P^ enzyme was obtained from Berkeley QB3 MacroLab. Frozen aliquots of H- Tn5^R27S,E54K,L372P^ enzyme stock (3.9mg/mL) suspended in Storage Buffer (50mM Tris pH 7.5, 800mM NaCl, 0.2mM EDTA, 2mM DTT, 10% glycerol) were thawed at 4°C, then diluted in Tn5 Dilution Buffer (50mM Tris pH 7.5, 200mM NaCl, 0.1mM EDTA, 2mM DTT, and 50% glycerol) to ∼1mg/mL Tn5 (18.9uM monomer) by rotational mixing at 4°C for 3.5h until fully homogenized. Tn5 was loaded with SMRT-Tag adaptors by gentle mixing of 1.02x volumes of 1mg/mL Tn5 with 1x volume of 20uM annealed SMRT-Tag adaptors using a wide-bore pipette, followed by an incubation at 23°C with continuous shaking at 350rpm for 55 min. Loaded Tn5 (SMRT-Tn5, 9.4uM monomer) can be supplemented with glycerol up to a final concentration of 50% and stored at −20° for up to 6 months.

### Assays for transposase activity

#### Confirming Tn5 Loading

Effective adaptor loading was confirmed by blue native PAGE gel-electrophoresis. Briefly, 1-2uL of SMRT-Tn5 combined with 4x Native Gel Loading Buffer (Invitrogen) was loaded per well on a NativePAGE 4-16% Bis-Tris Gel (Invitrogen) running at 150V for 1h at 4°C, followed by 180V for 15 min. Gels were stained with 1x SYBR Gold Solution (Invitrogen) in TAE, followed by 1x Coomassie Blue (Invitrogen) for 1h at room temperature, and imaged on an Odyssey XF imaging system (LI-COR).

#### Assaying Tagmentation Size Tunability

Tagmentation optimization was carried out in parallel using a serial dilution of stock SMRT-Tn5 in RNAse-free water. Diluted Tn5 was incubated with 160ng of genomic DNA (ProMega) using a range of buffers, temperatures and incubation times. Tagmentation reactions were terminated by addition of 0.2% SDS (final concentration 0.04%), and visualized via 0.4-0.6% 1X-TAE-Agarose gel. Electrophoresis run time was increased to 2-3h, and voltage decreased to 60-80V to maximize band resolution. Gels were stained with 1x SYBR Gold, and imaged on an Odyssey XF imaging system.

### Preparation of gDNA SMRT-Tag libraries

Purified High Molecular Weight genomic DNA (HG002-4, Coriell Institute) was normalized to 160ng per sample as input for SMRT-Tag library preparation, which included tagmentation, gap repair, exonuclease digestion and library validation.

For Tn5 tagmentation, reactions were prepared by adding Tagmentation Mix (10mM TAPS-NaOH pH 7.5, 5mM MgCl2, and 10% DMF) into each sample up to 9uL plus 1uL barcoded SMRT-Tn5 (0.073uM or 128x dilution of stock). Reactions were incubated at 55 °C for 30 min and terminated by an addition of 0.2% SDS (final concentration 0.04%) at RT for 5 min, followed by a 2x SPRI cleanup and elution in 12uL of 1x EB. For gap repair, tagmented samples were incubated in Repair Mix (0.1U Phusion-HF (New England Biolabs), 4U Taq DNA Ligase (New England Biolabs), 1x Taq DNA Ligase Reaction Buffer, 0.8mM dNTPs) at 37°C for 1 hr, followed by a 2x SPRI cleanup and elution in 12uL of 1x EB. To assess repair efficiency, 1uL of eluted sample post gap repair was measured by Qubit 1x High Sensitivity DNA Assay. For exonuclease digestion, reactions were incubated in ExoDigest Mix (100U Exonuclease III (New England Biolabs), 1x NEBuffer) at 37 °C for 1 hr, followed by a 2x SPRI cleanup and elution in 12uL of 1x EB. Libraries were multiplexed and pooled equimolarly based on the sample concentration measured by Qubit 1x High Sensitivity DNA Assay.

To optimize the loading efficiency, a size selection step using 35% (v/v) AMPure PB beads diluted in 1x EB was performed to enrich for libraries containing high molecular weight >5000 bp (HMW). 3.1x of 35% AMPure PB beads were added to the pooled library and the size-selected HMW was eluted in 15uL of 1x EB. Optionally, 0.25x AMpure PB beads were added to the supernatant and the low molecular weight <5000 bp (LMW) was eluted in 15uL of 1x EB.

To validate library quality, 1uL of eluted samples was run on Qubit 1x High Sensitivity DNA Assay and Agilent 2100 Bioanalyzer High Sensitivity DNA Assay to measure sample concentration and size distribution. HMW pool was sequenced on PacBio Sequel II 8M SMRTcells in-house using 2.2 polymerase, with LMW pool using 2.1 polymerase optionally. SMRT-Tag libraries were sequenced in-house over several pooled 30 h Sequel II movie runs with a 2 h pre-extension time and a 4 h immobilization time

### SAMOSA-Tag

#### Nuclei isolation

1-2 million OS152 cells were harvested by centrifugation (300xg, 4 °C, 10 min), washed in ice cold 1x PBS, and resuspended in 1 mL cold Nuclear Lysis Buffer (20 mM HEPES, 10 mM KCl, 1 mM MgCl2, 0.1% Triton X-100, 20% Glycerol, 1x Protease Inhibitor (Roche)) by gentle mixing with a wide-bore pipette. The suspension was incubated on ice for 5 min, then nuclei were pelleted (600xg, 4 °C, 10 min), washed with Buffer M (15 mM Tris-HCl pH 8.0, 15 mM NaCl, 60 mM KCl, 0.5 mM Spermidine), and counted via a Countess III cell counter (Thermo Fisher Scientific).

#### In situ SAMOSA footprinting

Permeabilized nuclei were pelleted (600xg, 4 °C, 10 min) and resuspended in 400uL Buffer M supplemented with 1 mM S-adenosyl-methionine (SAM, New England Biolabs) and 200uL aliquoted as a non-methylated control. Nonspecific adenine methyltransferase EcoGII (New England Biolabs, high concentration stock 25,000U/mL) was added to the reaction and incubated at 37 °C for 30 min with 300rpm shaking every 2 min. SAM was replenished to 1.16mM after 15 minutes in both the reaction and non-methylated control.

#### Tagmentation of footprinted nuclei

Methylated nuclei along with non-methylated controls were pelleted by centrifugation (600xg, 10 min) and gently resuspended in 250uL 1x Omni-ATAC Buffer (10mM Tris-HCl pH 7.5, 5mM MgCl2, 0.33x PBS, 10% DMF, 0.01% Digitonin (Thermo Fisher Scientific), 0.1% Tween-20). The nuclei suspension was then filtered through a 40µm cell strainer (Scienceware FlowMi), and aggregate dissociation was verified by counting and visualization via Countess III. Both methylated and non-methylated reactions were further distributed into 10,000 - 30,000 nuclei aliquots, and based on the desired fragment size post tagmentation, 9.4pmol - 37.6pmol of uniquely barcoded SMRT-Tn5 was added per reaction. Tagmentation reaction volumes were brought up to 50uL in 1x Omni-ATAC Buffer, then incubated at 55 °C for 1 hr.

#### Tagmentation termination and purification

To terminate tagmentation, reactions were pre-treated with 10uL of 10mg/mL RNase A (Thermo Fisher Scientific) at 37 °C for 15 min with 300rpm shaking. Termination Lysis Buffer (2.5uL of 20mg/mL Proteinase K (Ambion), 2.5uL of 10% SDS and 2.5uL of 0.5M EDTA) prepared at room temperature was added to the reaction, followed by an incubation at 60 °C with 1000rpm continuous shaking for at least 1h, up to 2h for improved lysis.

To extract tagmented fragments, 2x SPRI beads were added to the reaction, mixed until homogenous, and incubated at 23 °C for 30 min with mixing at 350 rpm every 3 minutes to keep the beads resuspended. Beads were pelleted via magnet, washed twice in 80% ethanol, then eluted in 20uL of elution buffer (EB, 10mM Tris-Cl pH 8.5) at 37 °C for 15 min with interval mixing at 350 rpm every 3 minutes to maximize sample recovery. Samples were subjected to an additional 0.6x SPRI cleanup to enrich for fragments > 500bp, and stored at 4°C overnight, or up to two weeks at -20°C.

#### Preparation of SAMOSA -Tag libraries

Purified, tagmented DNA extracted from OS152 nuclei (methylated and unmethylated control) were normalized to 160ng per sample as input for SAMOSA-Tag library preparation. A total of 8 methylated replicates along with 1 unmethylated control, each tagmented by a different set of barcoded SMRT-Tag adaptors, were processed in subsequent steps, including gap repair, exonuclease digestion and library validation. For gap repair, tagmented samples were incubated in Repair Mix (0.1U Phusion-HF, 4U Taq DNA Ligase, 1x Taq DNA Ligase Reaction Buffer, 0.8mM dNTP mix) at 37 °C for 1 hr, followed by a 2x SPRI cleanup and an elution in 12uL of 1x EB. To assess repair efficiency, 1uL of eluted sample post gap repair was measured by Qubit 1x High Sensitivity DNA Assay. For exonuclease digestion, reactions were incubated in ExoDigest Mix (100U Exonuclease III, 1x NEBuffer) at 37 °C for 1 hr, followed by a 2x SPRI cleanup and an elution in 12uL of 1x EB. To validate library quality, 1uL of eluted samples was run on Qubit 1x High Sensitivity DNA Assay and Agilent 2100 Bioanalyzer High Sensitivity DNA Assay to measure sample concentration and size distribution. Libraries were multiplexed and pooled equimolarly and sequenced on PacBio Sequel II 8M SMRTcells in-house using 2.1 polymerase. SAMOSA-Tag libraries were collected over several pooled 30 h Sequel II runs with a 2 h pre-extension time and a 4 h immobilization time.

### Data Analysis

All scripts and jupyter notebooks used for analyses are available at https://github.com/RamaniLab/SMRT-Tag. All plots were made using *ggplot2* (https://ggplot2.tidyverse.org/).

#### Data preprocessing

For all experimental data, HiFi reads were generated from raw subreads using *ccs* (v.6.4.0, Pacific Biosciences) with the additional flag *--hifi-kinetics* to annotate reads with kinetic information. *Lima* (v.2.6.0, Pacific Biosciences) with flag *--ccs* was used to demultiplex runs into sample-specific BAM files, and samples sequenced across multiple cells were merged using *pbmerge* (v1.0.0, Pacific Biosciences). Reads were aligned using *pbmm2* (v.1.9.0, Pacific Biosciences) to a modified version of the hg38 reference genome with the contaminant contig KI270752.1 removed, and read quality ascertained from predicted per-base QV scores and the estimated read quality generated by *ccs*.

#### SNV-based analysis of SMRT-Tag demultiplexing

The hs37d5 GRCh37 reference genome^39^, GIAB v4.2.1 benchmark^40^ VCF and BED files for HG002, HG003, and HG004, and GIAB v3.0 GRCh37 genome stratifications^25^ were accessed via the following links:

~~~
ftp://ftp-trace.ncbi.nlm.nih.gov/giab/ftp/release/references/GRCh37/hs37d5.fa.gz
ftp://ftp-trace.ncbi.nlm.nih.gov/giab/
ftp/release/AshkenazimTrio/HG002_NA24385_son/NISTv4.2.1/GRCh37/
ftp://ftp-trace.ncbi.nlm.nih.gov/giab/ftp/release/AshkenazimTrio/HG003_NA24149_father/NISTv4.2.1/GRCh37/
ftp://ftp-trace.ncbi.nlm.nih.gov/giab/ftp/release/AshkenazimTrio/HG004_NA24143_mother/NISTv4.2.1/GRCh37/
ftp://ftp-trace.ncbi.nlm.nih.gov/giab/ftp/release/genome-stratifications/v3.0/v3.0-stratifications-GRCh37.tar.gz
~~~

Private SNVs were for each individual were obtained using *bcftools* (v1.15.1) and regions for variant calling and evaluation comprising the union of the benchmark BED files were generated using *bedtools* (v2.3.0):

~~~
bcftools isec \
   --threads 4 \
   -n∼100 -w 1 \
   -c some \
   -Oz -o unique.HG002_GRCh37_1_22_v4.2.1_benchmark.vcf.gz \
   HG002_GRCh37_1_22_v4.2.1_benchmark.vcf.gz \
   HG003_GRCh37_1_22_v4.2.1_benchmark.vcf.gz \
   HG004_GRCh37_1_22_v4.2.1_benchmark.vcf.gz
bcftools isec \
   --threads 4 \
   -n∼010 -w 2 \
   -c some \
   -Oz -o unique.HG003_GRCh37_1_22_v4.2.1_benchmark.vcf.gz \
   HG002_GRCh37_1_22_v4.2.1_benchmark.vcf.gz \
   HG003_GRCh37_1_22_v4.2.1_benchmark.vcf.gz \
   HG004_GRCh37_1_22_v4.2.1_benchmark.vcf.gz
bcftools isec \
   --threads 4 \
   -n∼001 -w 3 \
   -c some \
   -Oz -o unique.HG004_GRCh37_1_22_v4.2.1_benchmark.vcf.gz \
   HG002_GRCh37_1_22_v4.2.1_benchmark.vcf.gz \
   HG003_GRCh37_1_22_v4.2.1_benchmark.vcf.gz \
   HG004_GRCh37_1_22_v4.2.1_benchmark.vcf.gz
cat HG002_GRCh37_1_22_v4.2.1_benchmark_noinconsistent.bed \
   HG003_GRCh37_1_22_v4.2.1_benchmark_noinconsistent.bed \
   HG004_GRCh37_1_22_v4.2.1_benchmark_noinconsistent.bed | \
sort -k1,1 -k2,2n -k3,3n | \
bedtools merge | \
bgzip | \
> HG002-4.calling_regions.bed.gz
~~~

Demultiplexed HG002, HG003, and HG004 SMRT-Tag were aligned to hs37d5 using the *minimap2* aligner (v2.15) and *pbmm2* (v1.9.0) and per-base coverage was tabulated using *mosdepth* (v0.3.3):

~~~
pbmm2 align \
   --log-level INFO \
   --log-file <OUTPUT_LOG> \
   --preset HiFi \
   --sort \
   --num-threads <THREADS> \
   --sample <SAMPLE_NAME> \ hs37d5.fa \
   <UNALIGNED_BAM> \
   <OUTPUT_BAM>
mosdepth \
   --threads <THREADS> \
   --use-median \
   --by GRCh37_notinalllowmapandsegdupregions.bed.gz \
   <OUTPUT_PREFIX> \
   <ALIGNED_BAM>
~~~

Given low depth of coverage, we naively called SNVs within regions defined in the GIAB benchmark BED files supported by at least 2 reads and with minimum mapping quality of 15 using *samtools mpileup* (v1.15.1) and a custom script.

~~~
samtools mpileup \
   --no-BAQ \
   --fasta-ref hs37d5.fa \
   --positions HG002-4.calling_regions.bed.gz \
   <ALIGNED_BAM> | \
bgzip > <OUTPUT_PLP_GZ>
zcat <OUTPUT_PLP_GZ> | \
plp2vcf.py -q <MIN_MAP_Q> -d <MIN_DEPTH> - | \
bgzip > <OUTPUT_VCF>
~~~

For each of HG002, HG003, and HG004, naive SNV calls were intersected with private benchmark SNVs in regions labeled ‘not difficult’ in the GIAB v3.0 genome stratification and covered by at least 2 SMRT-Tag reads using *bedtools* (v2.30.0), *samtools* (v1.15.1), and *bcftools* (v1.15.1). For example, the analysis for HG002 SMRT- Tag calls were intersected with HG003 benchmark private SNVs:

~~~
zcat HG002/mosdepth/HG002.per-base.bed.gz | \
awk -v D=2 ’{if ($4 >= D) print}’ | \
bedtools merge -i - | \
bedtools intersect \
   -u -a - -b GRCh37_notinalldifficultregions.bed.gz | \ bgzip \
> HG002.d2.GRCh37_notinalldifficultregions.bed.gz
bcftools isec \
   --threads <THREADS> \
   -n =2 -w 1 \
   -c some \
   --regions-file HG002.d2.GRCh37_notinalldifficultregions.bed.gz \
   -Oz -o HG002.q15.d2_vs_HG003_unique.vcf.gz \
HG002.q15.d2.vcf.gz \ unique.HG003_GRCh37_1_22_v4.2.1_benchmark.vcf.gz
bcftools index \
   -t -f \
   --threads <THREADS> \
HG002.q15.d2_vs_HG003_unique.vcf.gz
bcftools stats \
   --threads <THREADS> \ HG002.q15.d2_vs_HG003_unique.vcf.gz \
> HG002.q15.d2_vs_HG003_unique.stats
# Determine total number of covered SNVs:
bcftools view \
   --threads <THREADS> \
   --regions-file HG002.d2.GRCh37_notinalldifficultregions.bed.gz \
   -Oz -o HG002.d2_vs_HG003.base.vcf.gz \
   --types snps \ unique.HG003_GRCh37_1_22_v4.2.1_benchmark.vcf.gz
bcftools index \
   -t -f \
   --threads <THREADS> \ HG002.d2_vs_HG003.base.vcf.gz
bcftools stats \
   --threads <THREADS> \ HG002.d2_vs_HG003.base.vcf.gz \
> HG002.d2_vs_HG003.base.stats
~~~

#### HG002 small variant (SNV and indel) calling and benchmarking

In addition to the hs37d5 GRCh37 reference genome, GIAB v4.2.1 benchmark VCF and BED files for HG002, and GIAB GRCh37 v3.0 genome stratifications used in the genotype demultiplexing analysis, we downloaded publicly available HG002 PacBio Sequel II HiFi reads (SRX5527202), which were generated with ∼11 kb size selection and Sequel II chemistry 0.9 and SMRTLink 6.1 pre-release, and are available aligned to the same reference genome via GIAB:

~~~
ftp://ftp-trace.ncbi.nlm.nih.gov/giab/ftp/data/AshkenazimTrio/HG002_NA24385_son/PacBio_SequelII_CCS_11k b/HG002.SequelII.pbmm2.hs37d5.whatshap.haplotag.RTG.10x.trio.bam
~~~

We used the *minimap2* aligner and *pbmm2* for alignment of HG002 SMRT-Tag CCS reads to hs37d5 as before. Similarly, median total coverage for SMRT-Tag and GIAB PacBio reads was determined using *mosdepth*. Reads were subsampled to 3-, 5-, 10-, and 15-fold depths using *samtools* (v1.15.1) based on *mosdepth* median coverage:

~~~
samtools view \
   --threads <THREADS> \
   --subsample <FRAC>\
   --subsample-seed 0 \
   --bam \
   --with-header \
   --write-index \
   --output <OUTPUT_BAM> \
<ALIGNED_BAM>
~~~

Small variants (SNVs and indels) were called using *DeepVariant* (v1.4.0):

~~~
run_deepvariant \
   --model_type PACBIO \
   --num_shards <THREADS> \
   --verbosity 0 \
   --logging_dir <DIR> \
   --reads <ALIGNED_BAM> \
   --ref hs37d5.fa \
   --output_vcf <OUTPUT_VCF>
~~~

We then compared variants called from SMRT-Tag and HG002 PacBio Sequel II HiFi data against NIST v4.2.1 benchmarks^2^ using *hap.py* (v0.3.12) and GIAB v3.0 GRCh37 genome stratifications:

~~~
hap.py \
   -r hs37d5.fa \
   -o <OUTPUT_PREFIX> \
   -f HG002_GRCh37_1_22_v4.2.1_benchmark_noinconsistent.bed \
   --threads <THREADS> \
   --pass-only \
   --engine=vcfeval \
   --verbose \
   --logfile <OUTPUT_LOG> \
   --stratification v3.0-GRCh37-v4.2.1-stratifications.tsv \ HG002_GRCh37_1_22_v4.2.1_benchmark.vcf.gz \
   <DEEPVARIANT_VCF>
~~~

#### Predicting CpG methylation in single molecule reads

HiFi reads produced using both 2.1 and 2.2. sequencing polymerase chemistries were first demultiplexed with *lima* (v.2.6.0) to remove barcode sequences, then *primrose* (v.1.3.0) used to predict m^5^dC methylation status at CpG dinucleotides. Methylation probabilities encoded using the BAM tags ML and MM were parsed to continuous values and used for downstream single-molecule methylation predictions. Per-CpG methylation estimates were made using tools available at https://github.com/PacificBiosciences/pb-CpG-tools.

#### Predicting nucleosome footprints in SAMOSA-Tag data

SAMOSA-Tag data was preprocessed as above, and subsequently analyzed using a computational pipeline previously developed for detecting m^6^dA methylation in HiFi reads^31^. In brief, the kinetics of polymerase base addition were extracted per read, and a series of neural networks trained on methylated and unmethylated controls were used to predict the probability of m^6^dA methylation at all adenines on both the forward and reverse strands. Methylation probabilities were then binarized into accessibility calls using a two-state hidden Markov model. Accessibility information was encoded per read as a 0/1 modification probability using the BAM tags MM and ML for visualization using IGV.

#### U2OS CTCF ChIP-seq processing

Processed bed files from GEO accession GSE87831^41^ were lifted over from reference hg19 to hg38, and then analyzed as in Ramani *et al* (2019)^42^ to obtain predicted CTCF binding sites.

#### Insertion bias analyses at TSS and CTCF sites

Read-ends from SAMOSA-Tag data were extracted from BAM files and tabulated in a 5 kilobase window surrounding annotated GENCODEV28 (hg38) transcriptional start sites (TSSs) or ChIP-seq-backed CTCF binding sites. For visualization, all metaplots were smoothed with a running mean of 100 nucleotides. FRITSS / FRICBS was calculated as the fraction of read-ends falling within the 5 kilobase window.

#### CTCF CpG and accessibility analyses

m^6^dA accessibility signal surrounding predicted CTCF sites was extracted from accessibility pickle files and leiden clustered as in^31^. In addition to filtering out clusters that together accounted for less than 10% of all data, we also manually filtered out 1 cluster that corresponded to completely unmethylated fibers. Compared against all analyzed fibers surrounding CTCF sites, this cluster accounted for 3,627 fibers, or 11.5% of all CTCF-motif containing fibers. For CpG analyses, we used custom python scripts to convert CpG methylation to a similar pickle format as m^6^dA accessibility, and then used identical scripts to extract CpG methylation information per molecule, centered at CTCF sites. All data was then converted into text files for easy loading into R and visualization in ggplot2.

#### Classifying fibers by CpG content and CpG methylation

We binned all sequenced fibers by CpG content and CpG methylation to arrive at four bins, which we defined as high CpG content / methylation (*i.e.* > 0.5 average *primrose* score on a fiber; > 10 CpGs per kilobase), low CpG content / methylation (vice-versa), as well as high / low and low / high bins.

#### Fiber type clustering

We calculated single-molecule autocorrelograms and performed leiden clustering as in^31^. In addition to filtering out all clusters that together comprised less than 10% of all fibers, we also manually filtered out unmethylated / lowly methylated fibers, which fell out of the leiden clustering analysis and together accounted for 317,768 fibers (12.5% of all clustered fibers).

#### Fiber type enrichment

Fisher’s exact tests to determine fiber type enrichment were performed as in^31^.

## SUPPLEMENTARY FIGURES

**Supplementary Figure 1:**
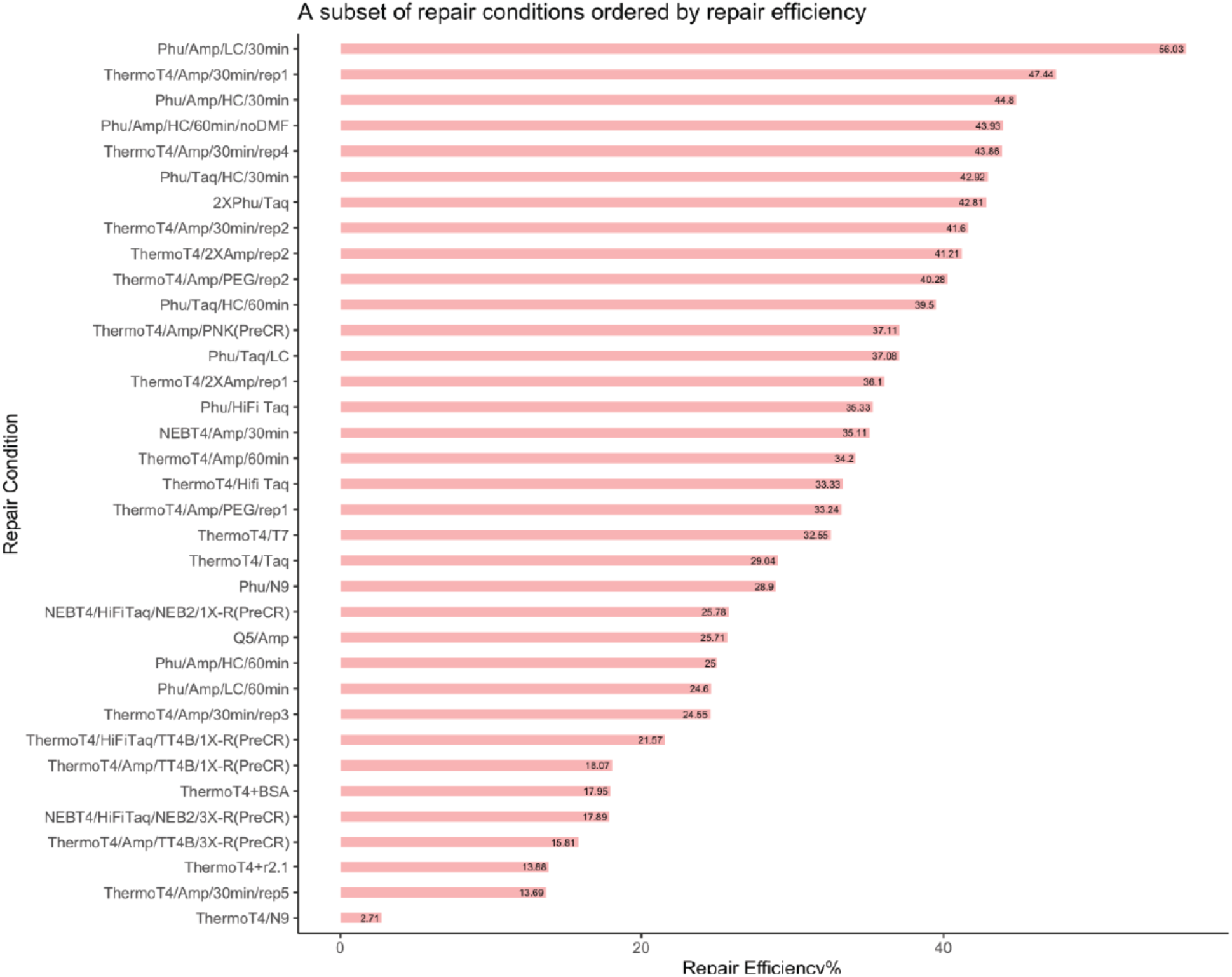
Tabulated repair efficiency for a subset of the 62 unique conditions tested to optimize gap repair. Repair efficiency (defined as the % yield of final product compared to input DNA by mass following exonuclease treatment) for 35 of the 62 conditions tested. We ultimately selected a mixture of Phusion polymerase and Taq ligase for gap repair as these provided the most consistently high repair efficiency across multiple experiments.

**Supplementary Figure 2.**
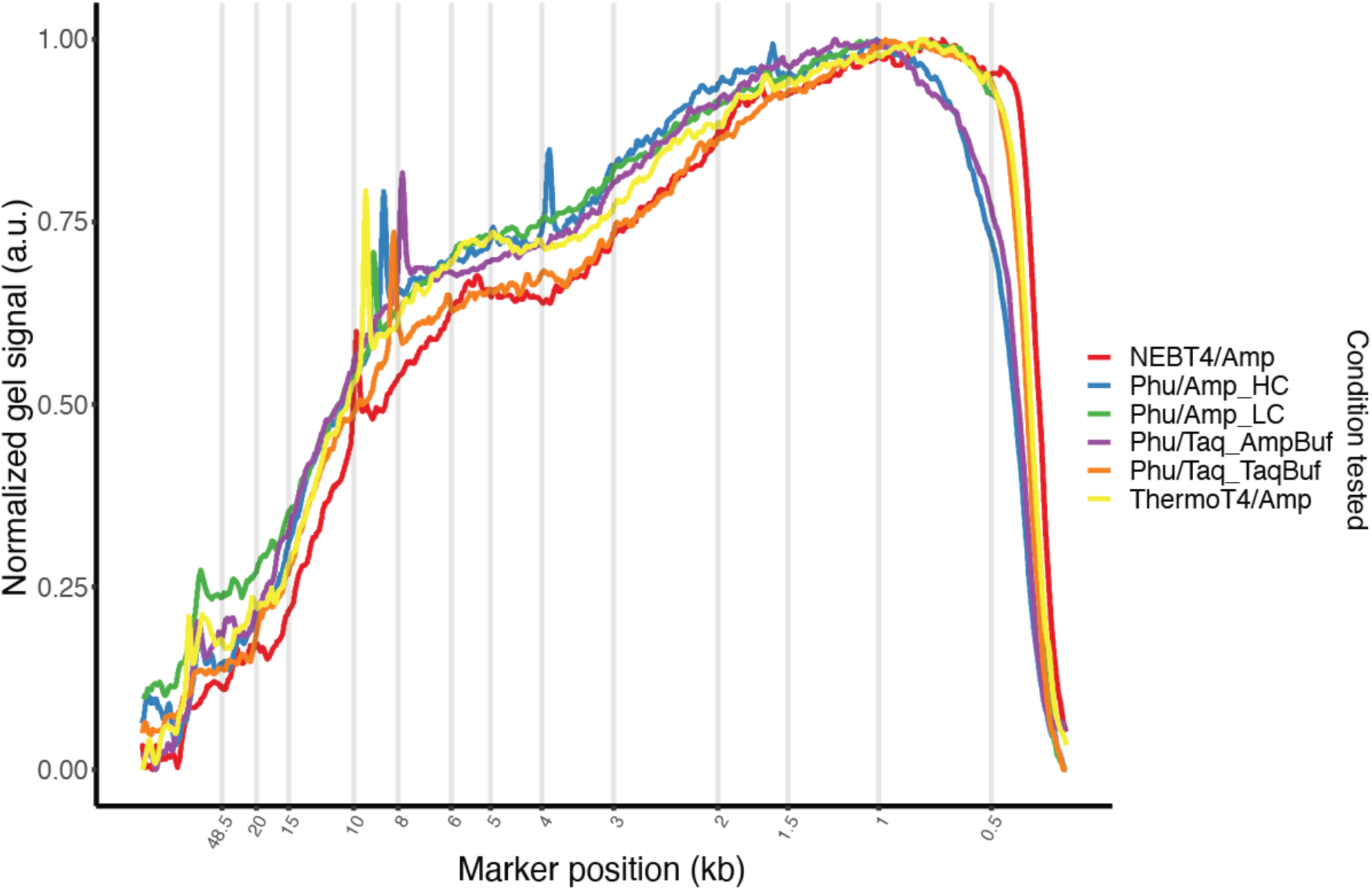
Example analytical gel trace for validating the size distribution of gap-repaired products for a subset of conditions. In addition to repair efficiency, we also validated that gap repair conditions did not appreciably change the size distribution of resulting libraries by gel electrophoresis. Shown here are analytical gel traces for six specific conditions tested in this study, including Phusion / Taq in multiple buffers.

**Supplementary Figure 3:**
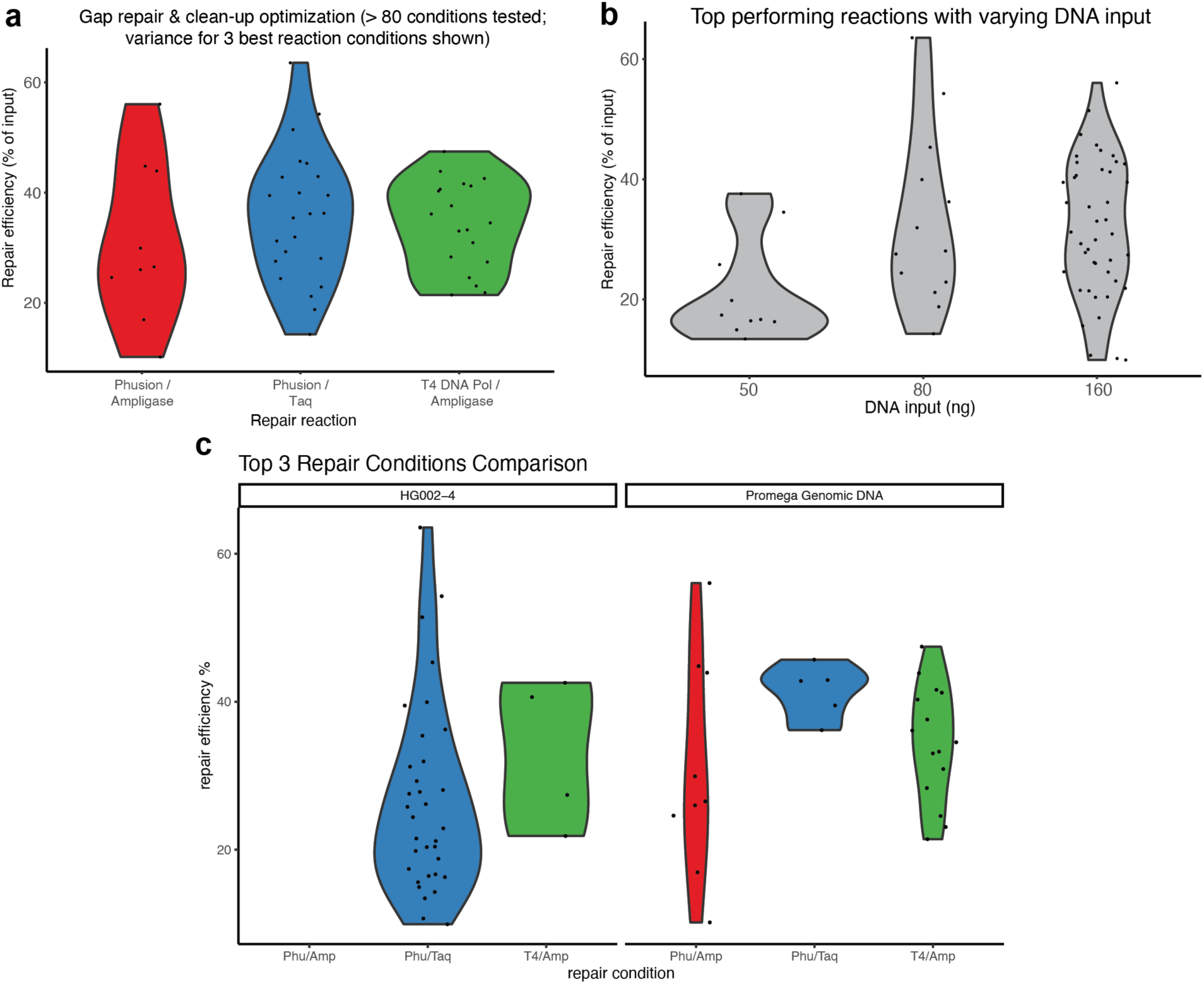
Optimizing gap repair conditions to maximize yield of closed SMRT-Tag reaction products. **a.)** A collection of repair efficiencies (measured as the % yield by mass of exonuclease-treated library compared to input DNA) for three different gap repair conditions. **b.)** Estimate of repair efficiency faceted by input DNA amount. We observed high repair efficiency down to 50 ng of input material. **c.)** Selection of Phusion / Taq versus T4 DNA polymerase / Ampligase repair conditions was motivated by increased reproducibility on high quality commercially available gDNA from Promega, compared to reactions with human samples HG002 – HG004, a subset of which were later determined to be slightly degraded.

**Supplementary Figure 4:**
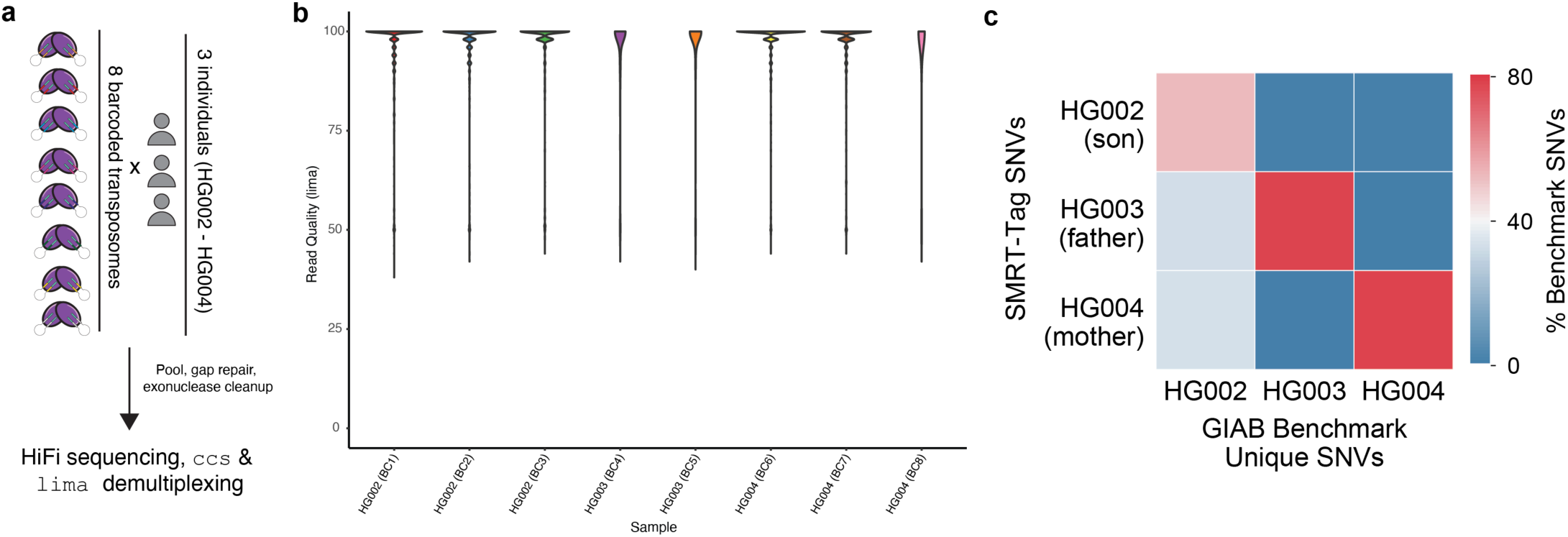
Control experiments to establish multiplexing with SMRT-Tag. **a.)** Schematic overview of a genotype mixing experiment where gDNA samples from HG003, HG004, and their progeny HG002 are individually barcoded with one of 8 different uniquely-loaded transposomes, pooled, gap-repaired, and sequenced on the PacBio Sequel II. **b.)** Demultiplexing results from PacBio’s proprietary *lima* barcode splitting software, demonstrating minimal cross-contamination across transposome barcodes / samples. **c.)** Percentage shared genotype information across barcoded samples. As expected HG002 harbors shared SNPs with HG003 and HG004, but HG003 and HG004 samples have minimal shared genotype overlap. For this analysis, all ‘private’ SNVs across HG003 and HG004 were considered.

**Supplementary Figure 5.**
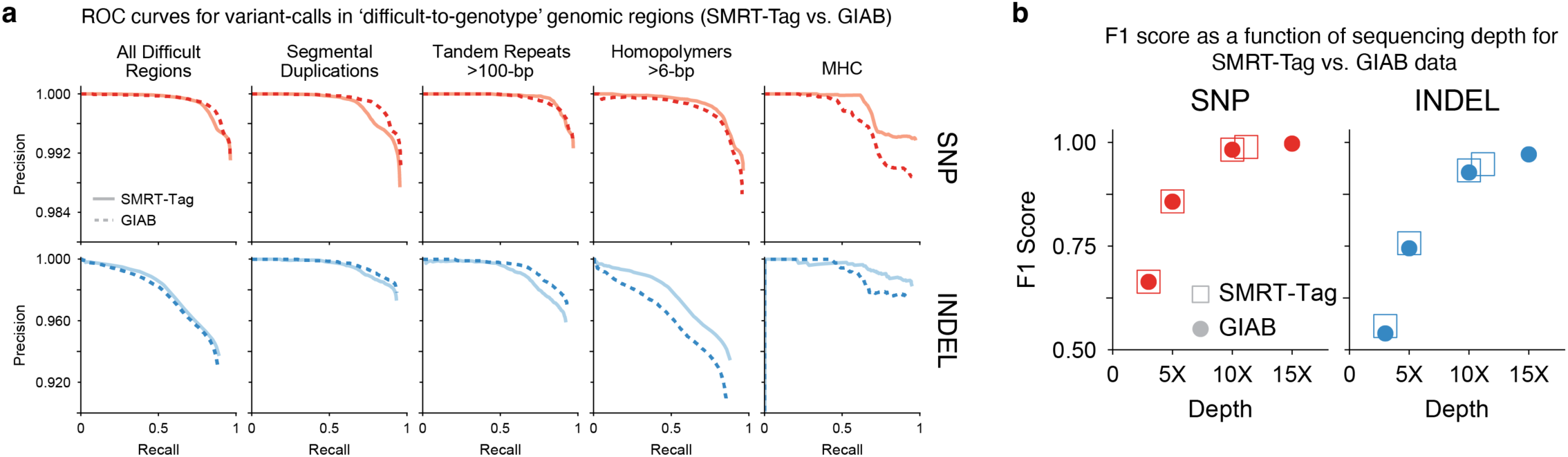
Genotyping performance of SMRT-Tag data across difficult-to-genotype regions and as a function of sequencing depth. **a.)** DeepVariant precision / recall curves for SNP (red) and indel (blue) variant calls in challenging genomic regions, including segmental duplications, tandem repeats, homopolymers, and the MHC locus, for SMRT-Tag data (solid) versus GIAB data (dashed). **b.)** Composite F1 score for SMRT- Tag (closed circles) versus GIAB data (open square) as a function of sequencing depth, for SNP (red) and indel (blue) variant calls.

**Supplementary Figure 6.**
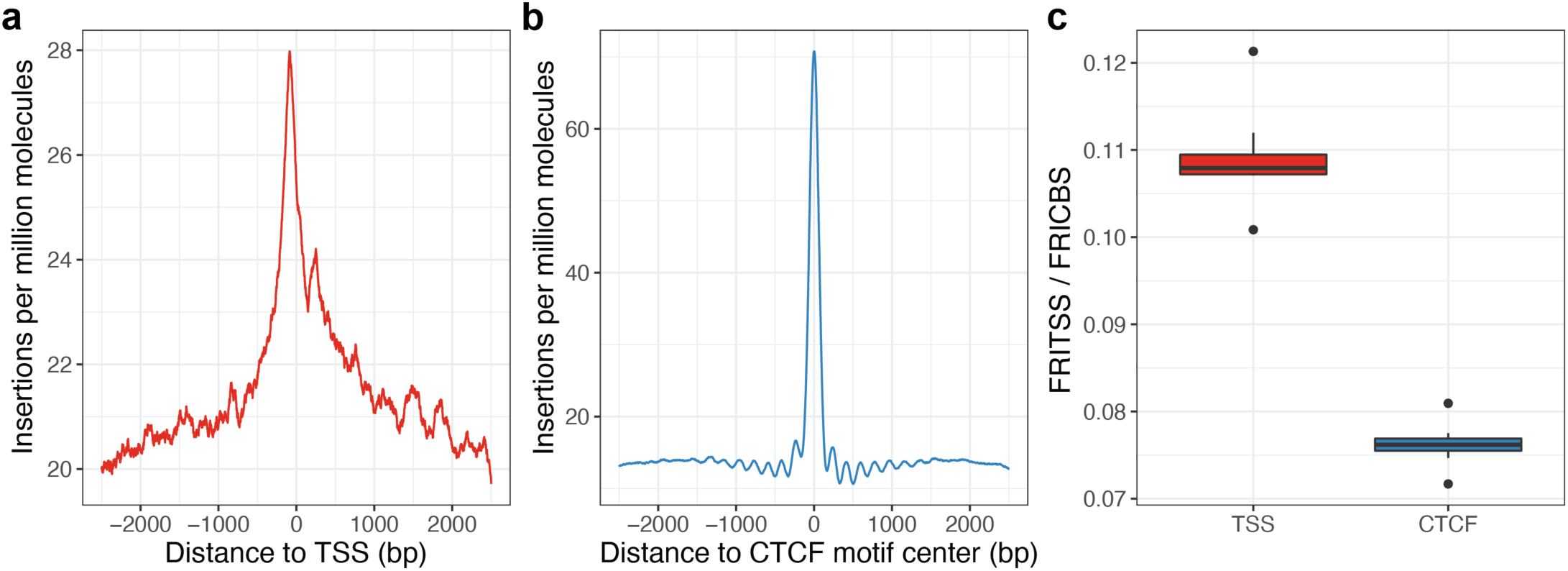
SAMOSA-Tag libraries demonstrate slight insertional bias at transcription start sites and CTCF motifs. **a.)** Metaplot of insertions per million sequenced molecules at hg38 transcriptional start sites (TSSs), in a 5 kb window centered at the TSS. Signal was smoothed using a 100 nt running mean. **b.)** Metaplot of insertions per million sequenced molecules at U2OS ChIP-seq backed CTCF binding sites, in a 5 kb window centered at the center of the CTCF motif. Signal was smoothed using a 100 nt running mean. **c.)** Boxplots of fraction of insertions in TSS (FRITSS) and fraction of insertions in CTCF binding sites (FRICBS) across all eight replicate experiments.

**Supplementary Figure 7.**
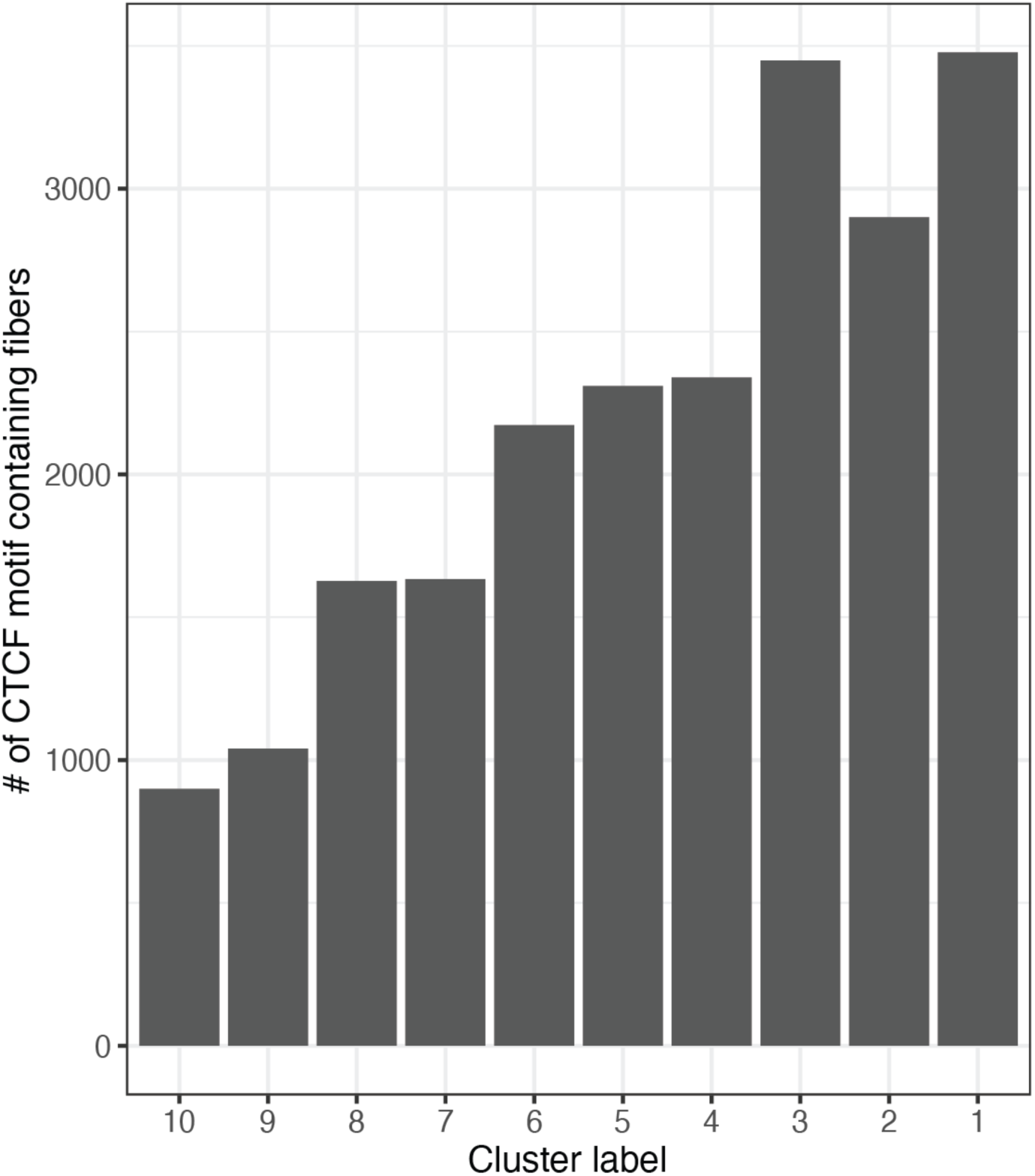
Cluster sizes resulting from leiden clustering of single-molecule accessibility patterns surrounding predicted CTCF sites. Cluster labels match the labels used in Figure 4B**,C**.

**Supplementary Figure 8.**
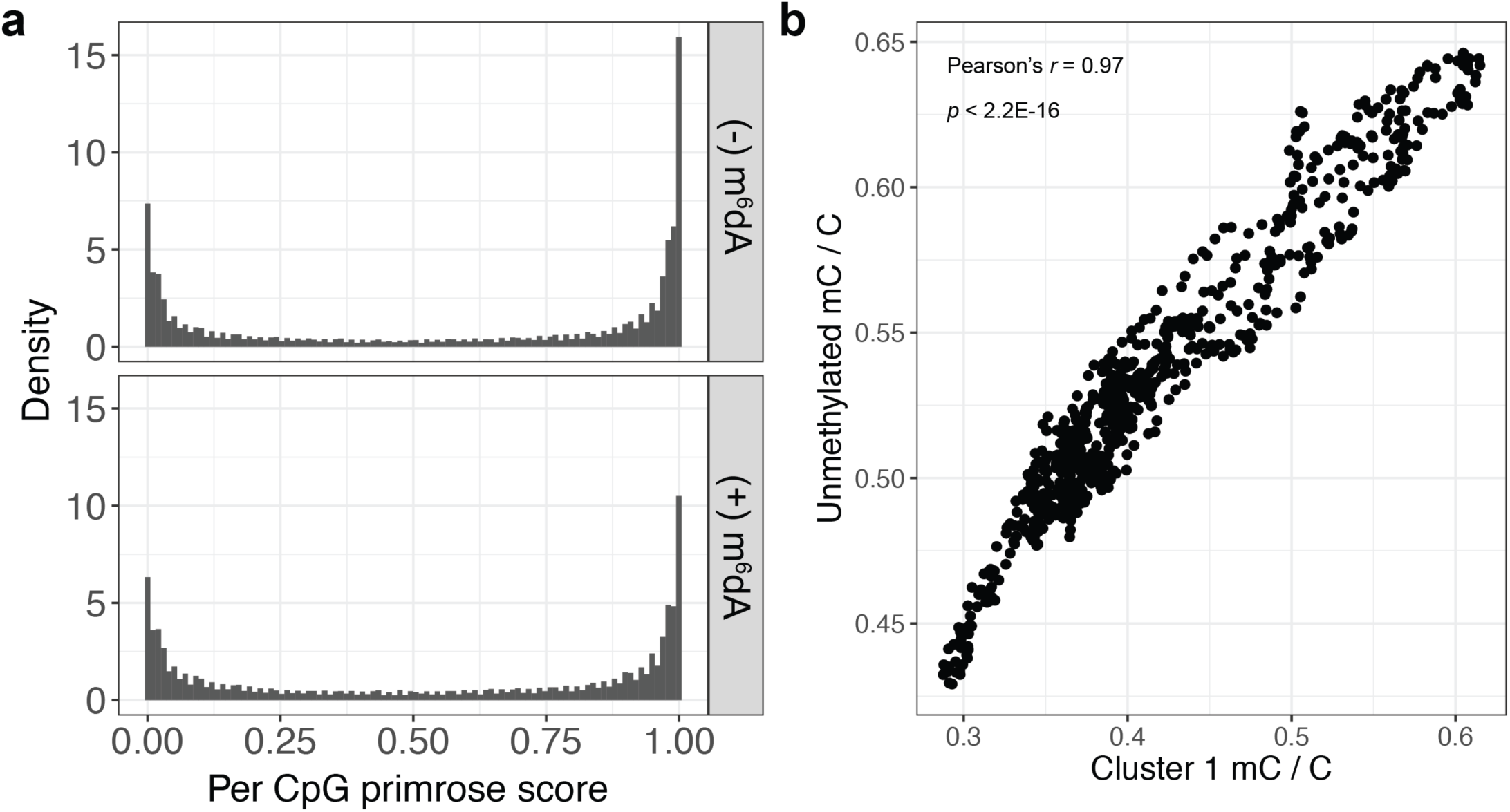
m^6^dA footprinting does not appreciably impact primrose CpG methylation predictions. **a.)** Distribution of per CpG *primrose* scores (50,000 sampled CpGs per experiment) for SAMOSA- Tag control experiments where EcoGII was omitted (top) and SAMOSA-Tag experiments (bottom). **b**.) Correlation of averaged CpG methylation signal from SAMOSA-Tag molecules without any detectable m^6^dA methylation surrounding predicted CTCF sites, versus CpG methylation from SAMOSA-Tag molecules from occupancy cluster 1. Signals were correlated with Pearson’s *r* of 0.97 (*p* < 2.2E-16).

**Supplementary Figure 9:**
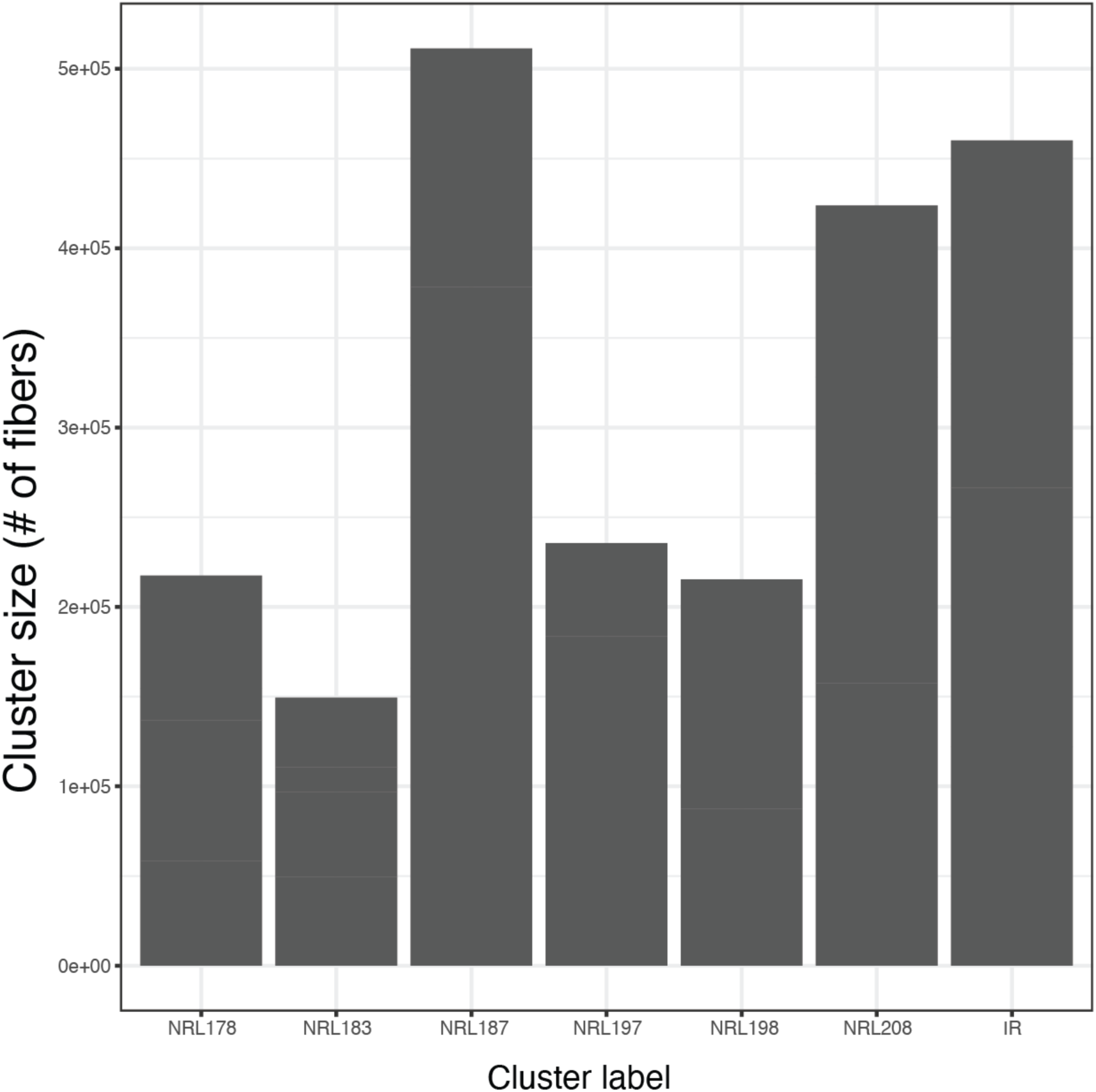
Cluster sizes resulting from leiden clustering of single-molecule autocorrelograms. Cluster labels match labels used in Figure 4E**,F**.

**Supplementary Figure 10.**
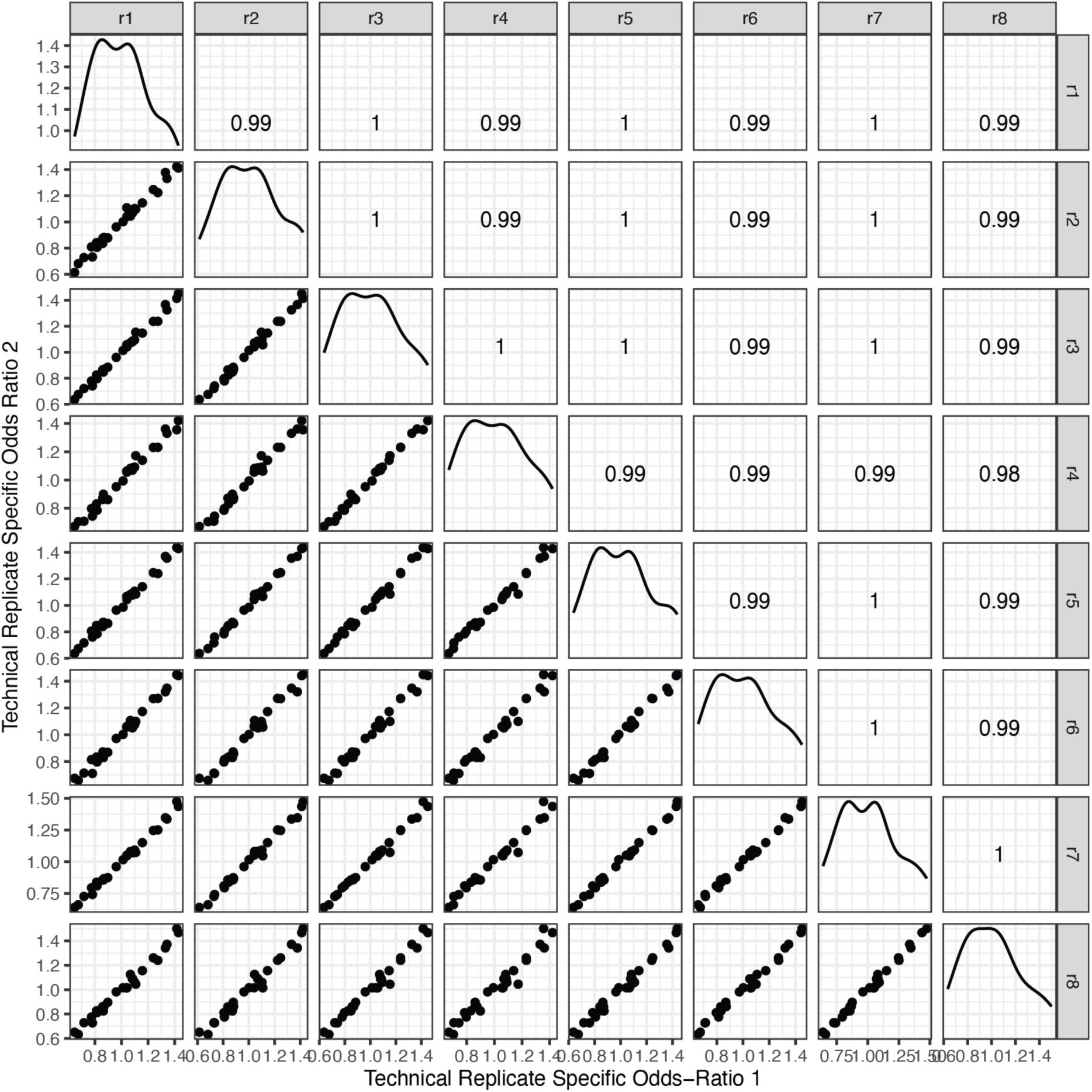
SAMOSA-Tag fiber enrichments in different CpG content / CpG methylation bins are technically reproducible. Matrix of scatter plots plus Pearson’s *r* correlation values across each of eight different replicate OS152 SAMOSA-Tag experiments.

## SUPPLEMENTARY TABLES

**Supplementary Table 1.**
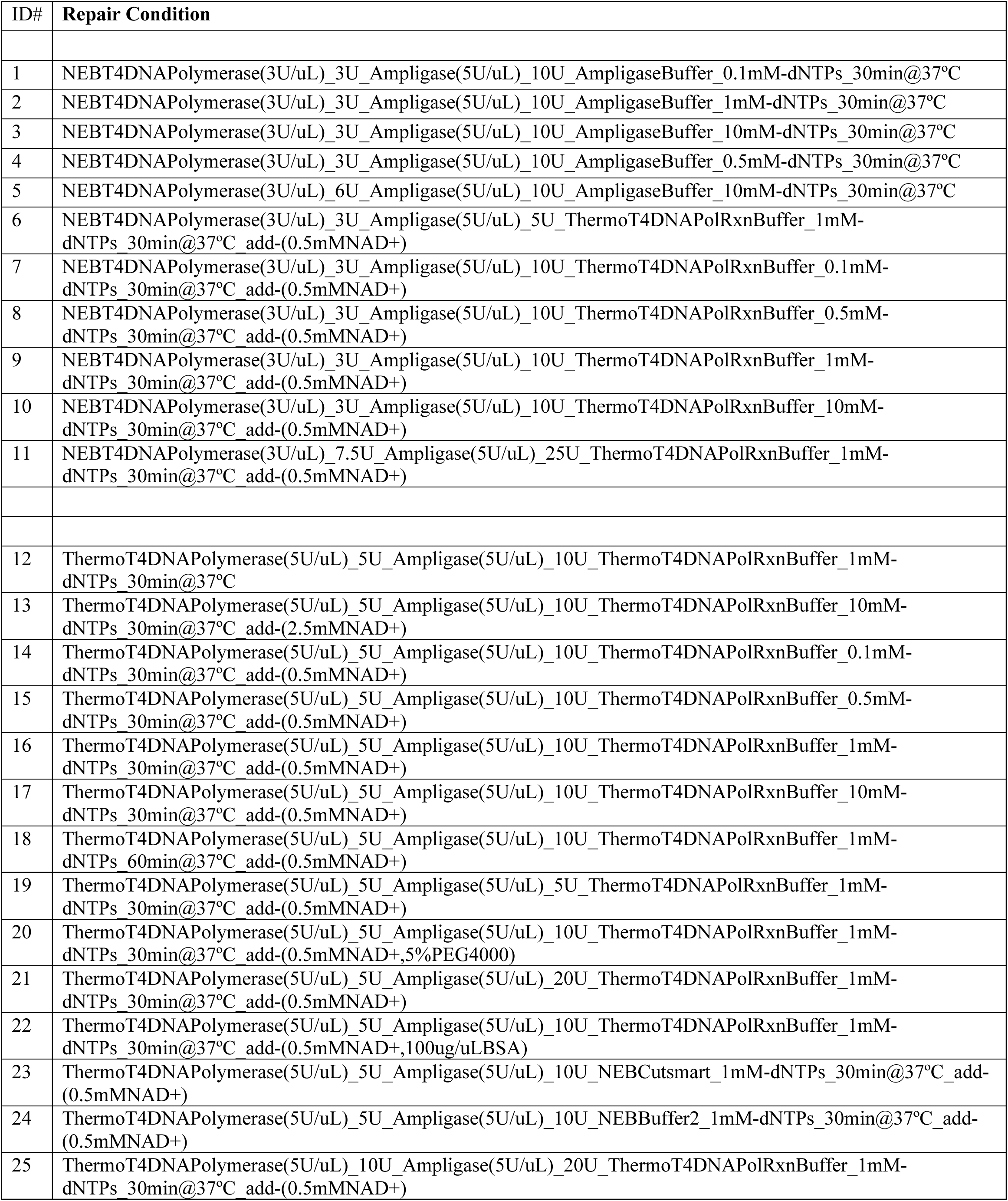
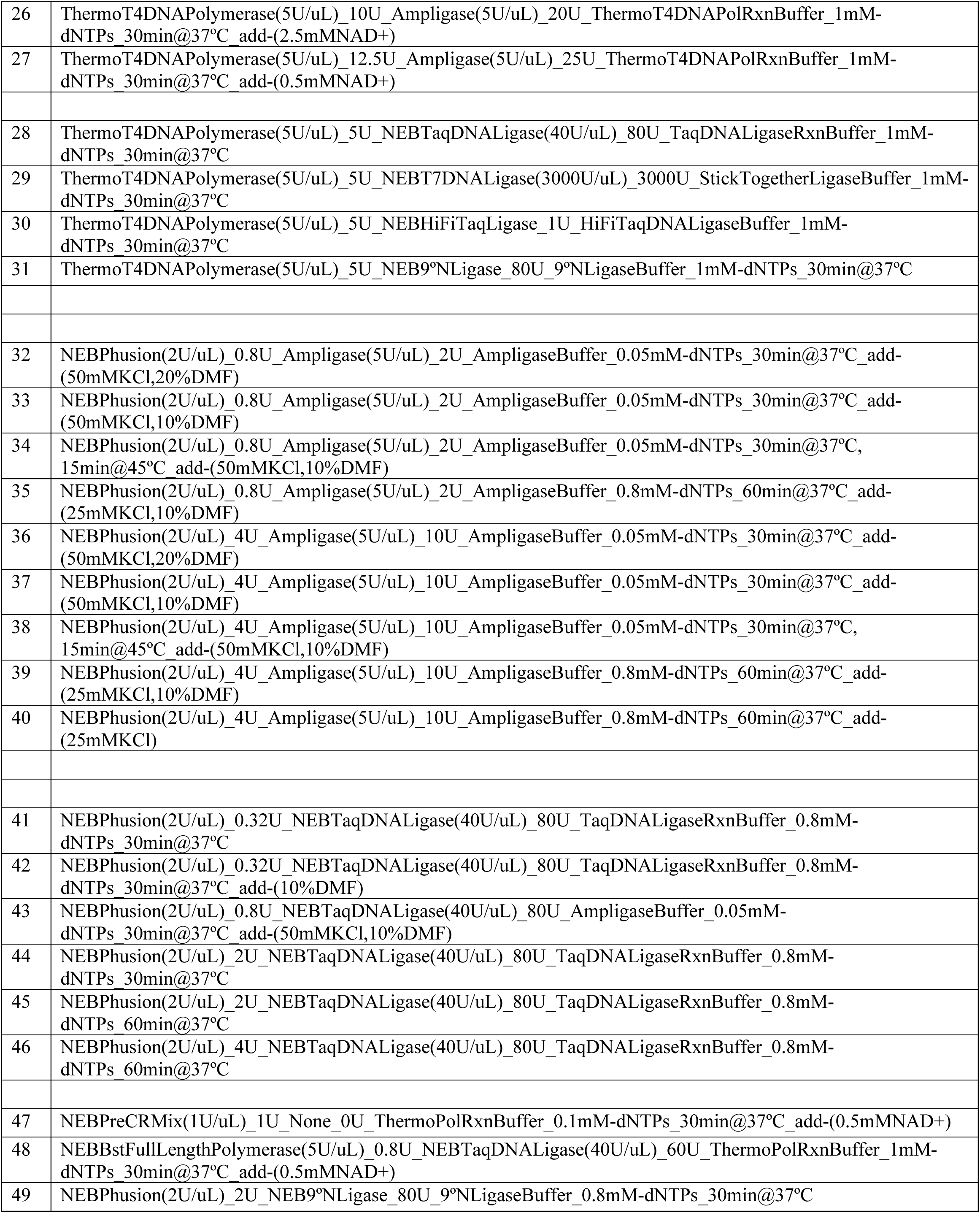
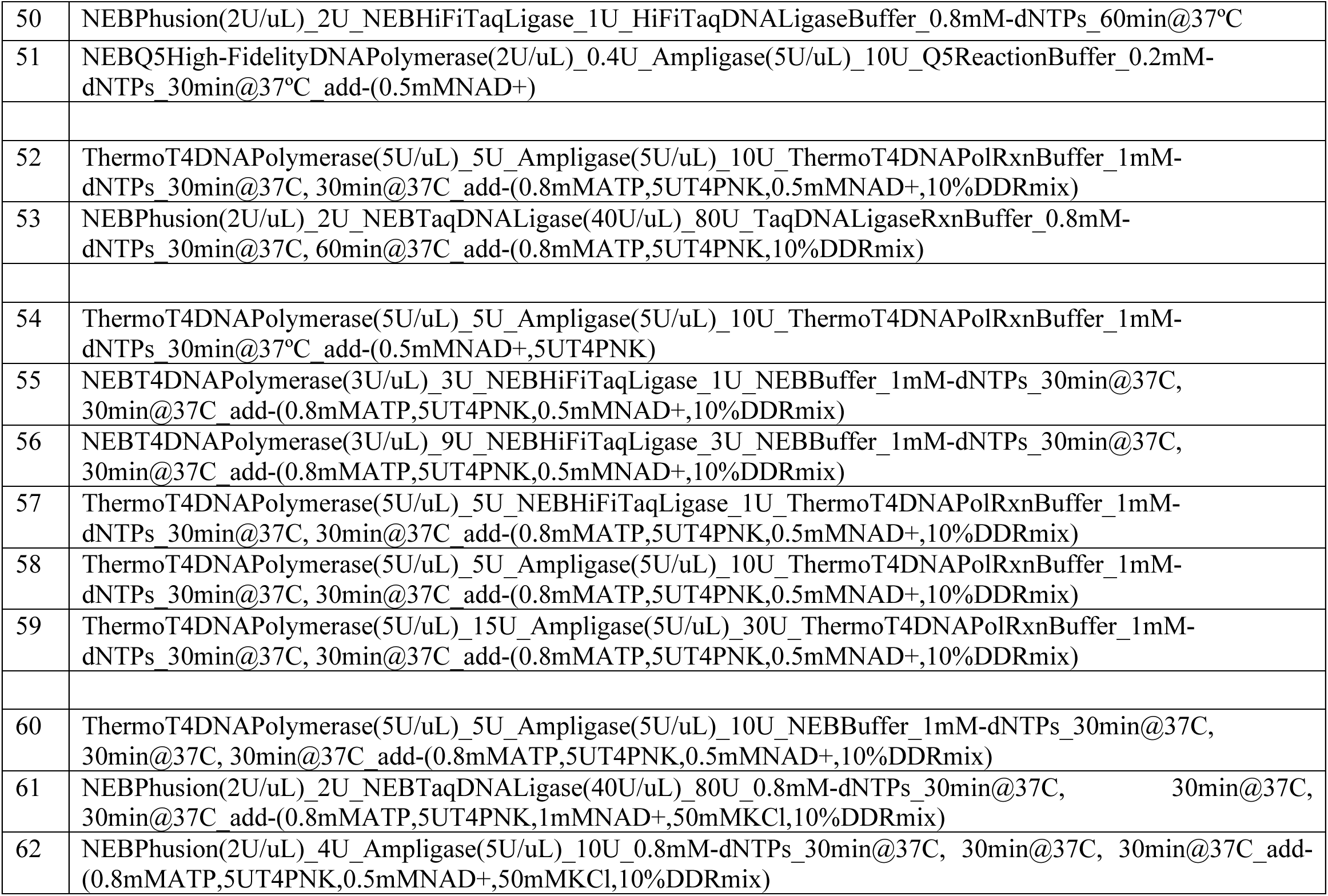
Summary of 62 gap repair conditions tested in this study.

**Supplementary Table 2:**
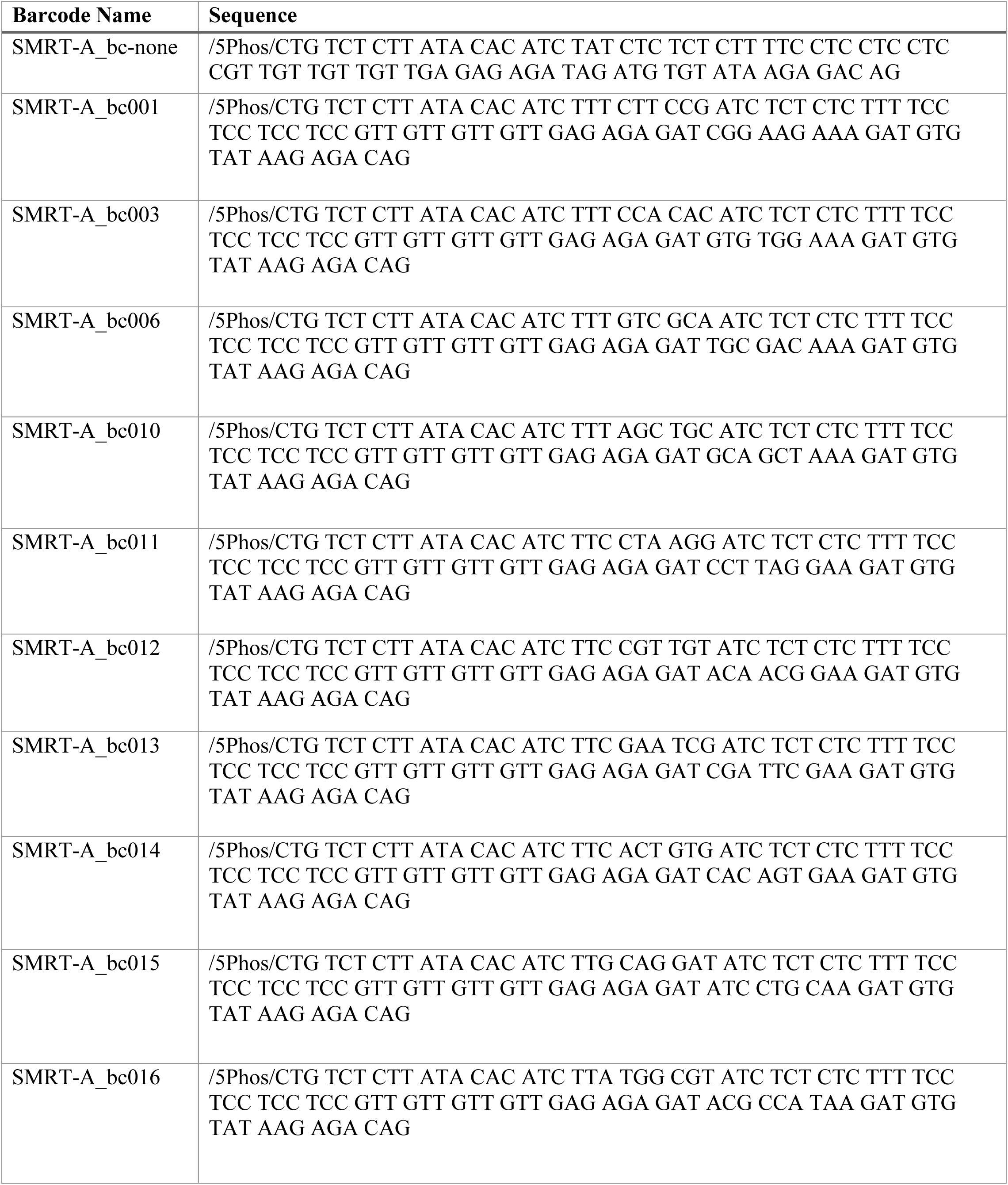
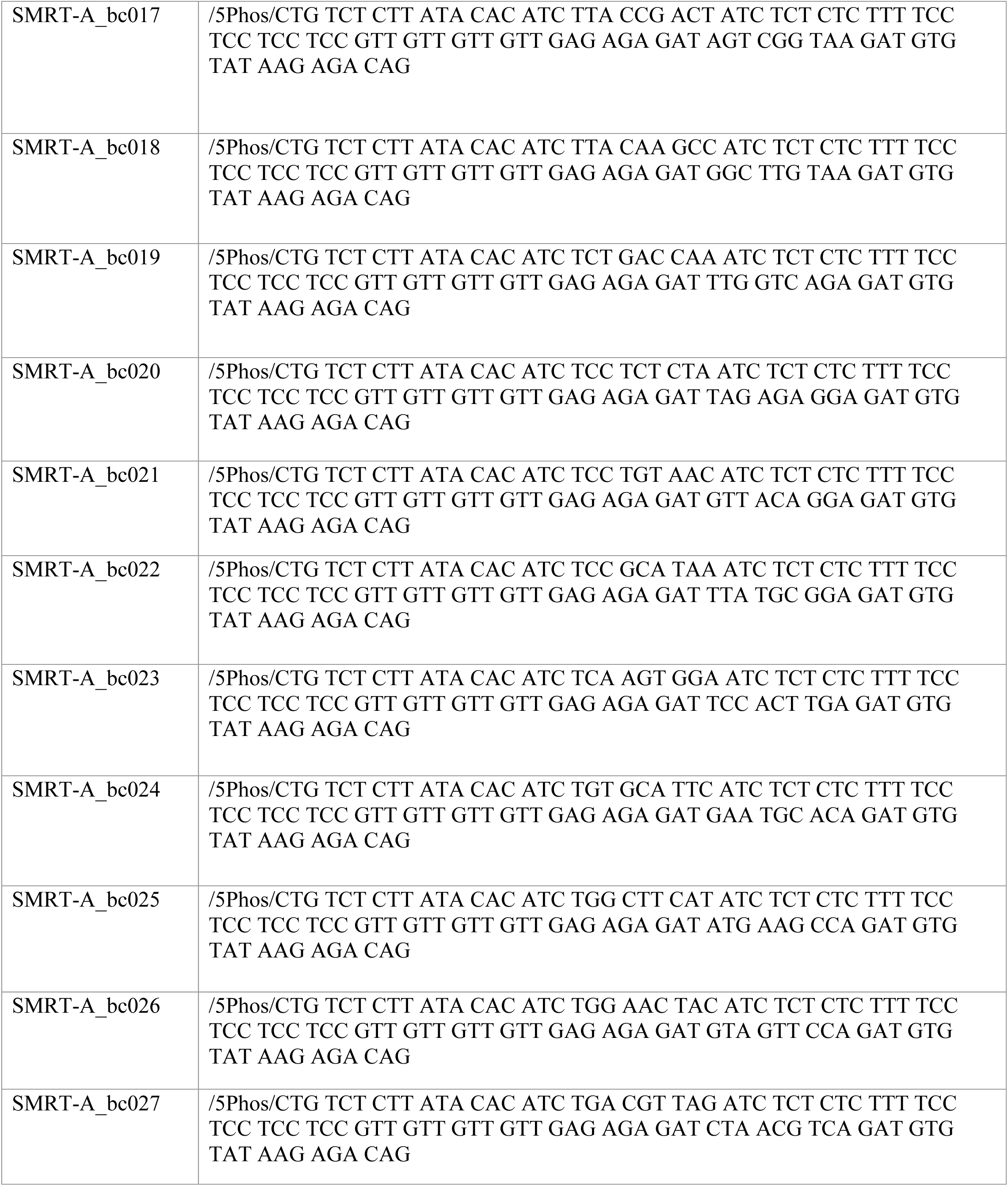
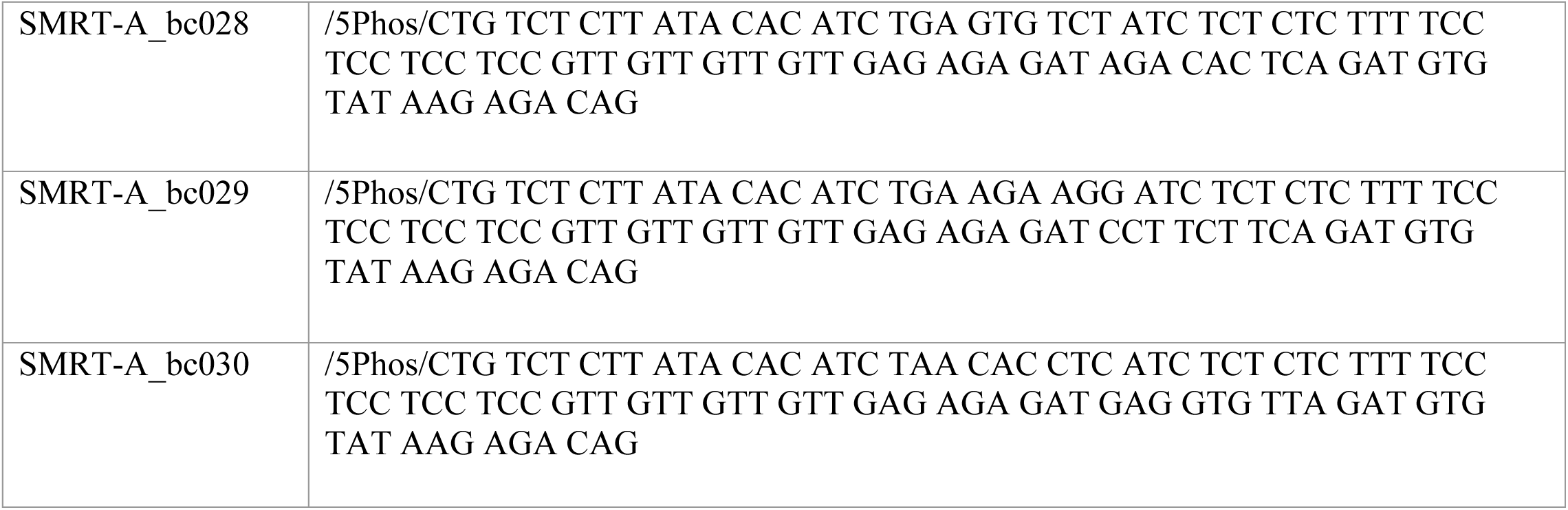
Barcoded SMRT-Tag adaptor sequences.

